# Deacetylation of β-Mannans by Two Complementary Carbohydrate Esterases from the Human Gut Microbe *Bacteroides cellulosilyticus*

**DOI:** 10.64898/2025.12.12.693890

**Authors:** Lars Jordhøy Lindstad, Pascal Michael Mrozek, Gordon Jacob Boehlich, Shaun Leivers, Phillip B. Pope, Sabina Leanti La Rosa, Bjørge Westereng

## Abstract

β-Mannans are widespread in the human diet as components of plant-derived foods and as food additives. Several classes of β-mannans are decorated with acetylations, which are key for their physicochemical properties and protection against enzymatic degradation. While the process for depolymerization of acetylated β-mannans has been described in depth for members of the phylum Bacillota, there is limited mechanistic knowledge on how Bacteroidota utilize these glycans. Here, we combined proteomics and biochemical analyses to functionally characterize a pair of carbohydrate esterases (CEs) from *Bacteroides cellulosilyticus* that, together, deacetylate complex β-mannans. We demonstrate that the newly identified *Bc*CExxx enzyme, representing a novel carbohydrate esterase (CE) family, exhibits high specificity by selectively removing axially oriented 2-*O*-acetyl groups from mannose residues. In contrast, *Bc*CE7 functions as a broad-spectrum esterase, capable of deacetylating oligosaccharides derived from structurally diverse substrates, including β-mannans, xylans, and acetylated cellulose. In transesterification reactions, *Bc*CExxx showed activity on both mannooligosaccharides and polymeric glucomannan. Overall, our findings provide new insight into the strategies that beneficial *Bacteroides* have evolved to deacetylate complex β-mannans in the human gut.

**Significance Statement:** β-Mannans, commonly found in various plant-based foods and used as food additives, can be metabolized by gut microbes, potentially impacting host health. Acetylation of β-mannans enhances their resistance to enzymatic breakdown, making carbohydrate esterases crucial for gut bacteria to utilize these carbohydrates. This study explores the roles of two carbohydrate esterases (CEs) from a human commensal *Bacteroides* species in deacetylating complex β-mannans. The two esterases together remove acetylations at different positions on mannose units. *Bc*CE7 is a versatile esterase that deacetylates a variety of oligosaccharides, while the newly discovered *Bc*CExxx is highly active towards 2-*O*-acetylations on mannose units in both oligosaccharides and polymeric β-mannan. This research enhances our understanding of how Bacteroidota species deacetylate β-mannans.

## Introduction

β-Mannans are hemicelluloses found in many edible plants and are used as plant-based food additives for their stabilizing, thickening, and gelling properties (1–4). They are widespread in important food sources like the endosperm of legumes, coffee beans, tomato seeds, bananas, seeds from monocots (corn, wheat, rice), and flax seeds, and contribute to the structure of plant cell walls (5–7). Depending on the source, β-mannan structures are classified into four substructures: linear mannan, galactomannan, glucomannan, and galactoglucomannan (Fig. 1a). The β-mannan backbone consists of β-1,4-linked D-mannopyranose moieties in linear mannan, which can be interspersed with β-1,4-linked D-glucopyranose moieties (glucomannan), and decorated with α-1,6-linked D-galactose moieties (galactomannan and galactoglucomannan) (8). In addition, β-mannans can be substituted to varying degrees, with acetylations on the D-mannopyranose moiety at the 2-*O*, 3-*O*, and 6-*O* positions (9). Acetylated galactoglucomannan (AcGGM) is common in the β-mannan from softwood, such as Norway spruce (*Picea abies*), where β-mannan accounts for about 20 % of the dry wood (10). Although the presence of acetyl groups in some of the sources described above is reported, documentation on acetylations remains limited. This is possibly due to alkali isolation procedures, which are commonly used, that readily remove acetylations from the carbohydrate. As many plant sources are found to express mannan *O*-acetyltransferases (also referred to as MOATs), it is suggested that the importance of acetylations is still underrepresented (11). Notably, the axially oriented 2-*O*-acetylation is structurally different from other common dietary (poly-/oligo-) saccharides, which have acetylations oriented in the equatorial plane (12). These features give β-mannans, such as in *Amorphophallus konjac*, guar gum, and locust bean gum, different physicochemical properties as food additives, while the degree of acetylation affects their solubility and ability to form gels (2, 3, 13, 14).

**Fig. 1.**
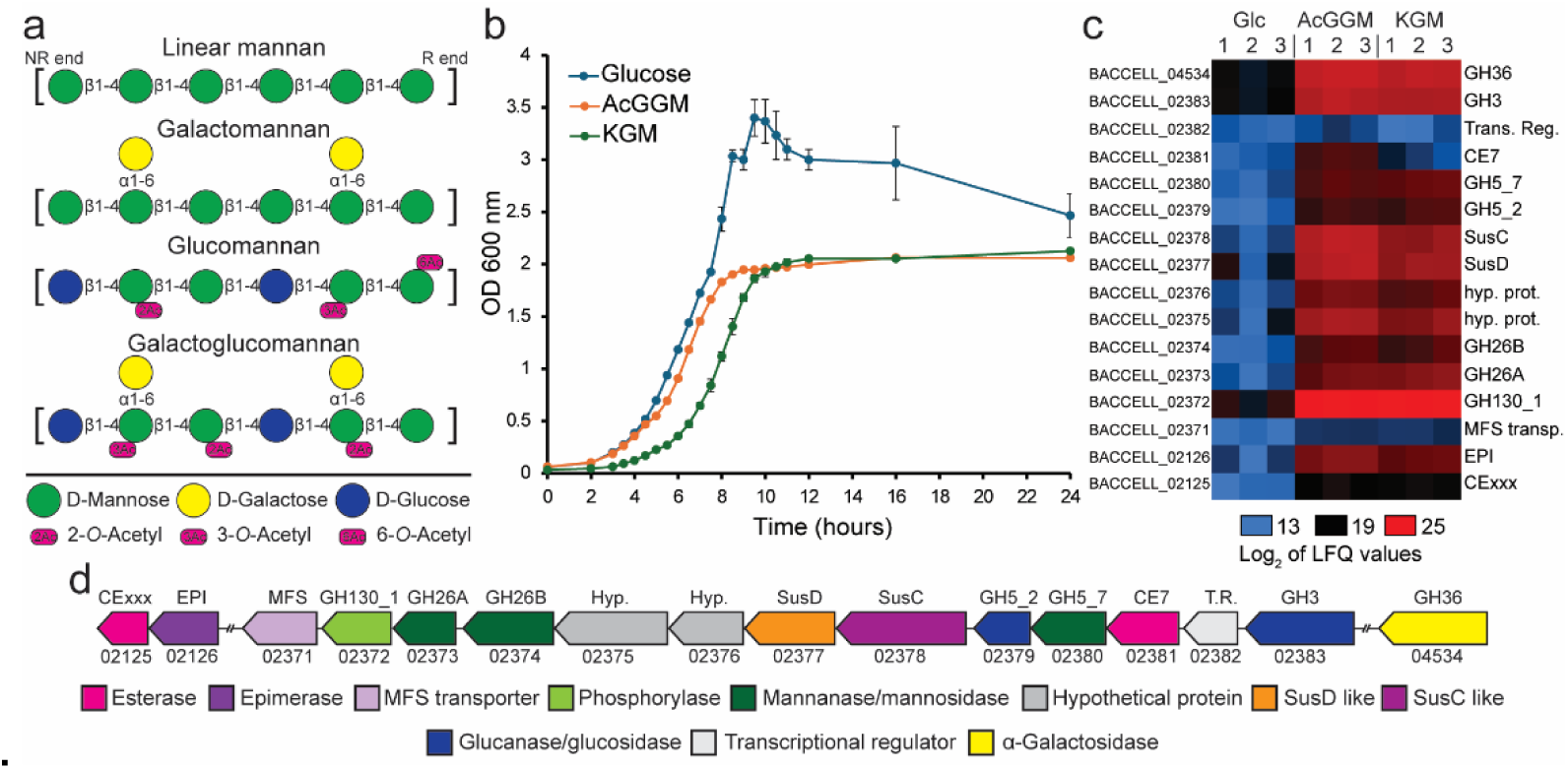
a) Various structures of β-mannans found in plants. Structural differences and degree of acetylation vary between sources as well as the processing of material for β-mannan isolation. NR end, non-reducing end; R end, reducing end. b) Growth of *B. cellulosilyticus* in minimal media supplemented with 5 mg/mL glucose, AcGGM, or KGM as the sole carbon source. c) The heat map presents the more abundant proteins from *B. cellulosilyticus* when grown on AcGGM and KGM compared to glucose. Each substrate was tested in triplicate, and the color intensity shows the protein abundance as Log_2_ of the LFQ values (13 to 25, in blue and red, respectively). d) Genomic representation of the MUL (BACCELL_02371-02383) with three additional genes (BACCELL_02125-02126 and BACCELL_04534) involved in β-mannan processing (based on proteomics data).

Dietary fibers, including β-mannans, can be depolymerized and further fermented by the gut microbiota into short-chain fatty acids (SCFAs), which are known to have beneficial effects on host health (15–19). Several bacterial species in the human gastrointestinal tract have been demonstrated to utilize β-mannans. In an in vitro study, members of the genera *Lactobacillus*, *Bifidobacterium*, and *Bacteroides* utilized Norway spruce-derived AcGGM as the sole carbon source (20). *Roseburia hominis* and *Bifidobacterium adolescentis* have been shown to grow on mannooligosaccharides obtained from galactomannan and AcGGM (21). Spruce AcGGM has also been shown to stimulate the growth of beneficial bacteria in the pig gut microbiota (22).

The structure of complex β-mannans requires the coordinated activity of various Carbohydrate-Active Enzymes (CAZymes), such as glycoside hydrolases (GHs) and carbohydrate esterases (CEs), for the complete depolymerization of this glycan into monosaccharides. The degradation apparatus for β-mannans has been described for *Roseburia intestinalis*, a butyrate-producing bacterium prevalent in the healthy gut (23). *R. intestinalis* L1-82 contains two Polysaccharide Utilization Loci (PUL) for β-mannan degradation, hereafter referred to as Mannan Utilization Loci (MUL), which were upregulated during growth on konjac glucomannan (KGM) and spruce AcGGM. The MULs contain an extracellular endomannanase (*Ri*GH26) that cleaves polymeric β-mannans into mannooligosaccharides, which are then transported into the cell and further degraded into monosaccharides. Another prominent member of the Bacillota phylum in the healthy gut microbiota, *Faecalibacterium prausnitzii*, harbors MULs similar to *R. intestinalis* (24). *F. prausnitzii*, however, lacks the gene coding for an extracellular endomannanase but can cross-feed on mannooligosaccharides derived from the mannanolytic activity of other bacteria. The MULs of both *R. intestinalis* and *F. prausnitzii* have two acetyl esterases that, together, have been demonstrated to deacetylate complex β-mannans completely (24, 25).

*Bacteroides* (Bacteroidota) species are abundant in the human gut and are known for their ability to utilize a broad range of dietary glycans (26, 27). Among them, *B. cellulosilyticus* has been shown to allocate a significant part of its genome (10%) for genes encoding CAZymes, conferring exceptional saccharolytic capabilities (28). In addition, it has been shown to provide beneficial health effects and attenuate experimental colitis in mice by producing zwitterionic capsular polysaccharides that influence T regulatory cells (29). Currently, some studies have demonstrated the growth of *Bacteroides* on galactomannan and glucomannan, although their mannan degradation systems are only partially characterized. Martens et al. showed that *B. ovatus* upregulates a MUL when grown on galactomannan and glucomannan (30), and the degradation apparatus includes two GH26 mannanases (*Bo*Man26A and *Bo*Man26B) and a GH36 α-galactosidase (*Bo*Gal36A) (31, 32). *Bo*Man26B is a surface-exposed endo-acting mannanase, while *Bo*Man26A generates mannobiose in the periplasm. *Phocaeicola dorei* (formerly known as *B. dorei*) contains a MUL that includes a gene encoding a GH26, which was highly upregulated when the bacterium was grown on either carob galactomannan (CGM) or KGM (33). *B. fragilis* possesses a GH26, characterized as a mannobiose-producing mannanase, along with an epimerase and mannosylglucose phosphorylase (34, 35). Finally, a study by McNulty et al. showed that *B. cellulosilyticus* has a PUL that is expressed when grown on galactomannan and glucomannan (28).

The complete degradation of acetylated β-mannans (e.g., *Aloe vera* mannan, KGM, and spruce mannan) requires esterases to remove acetylations as they contribute to steric hindrance and limit the activity of other enzymes (9, 36). Currently, there are 20 (1-9 and 11-21) CE families in the CAZy database (http://www.cazy.org/Carbohydrate-Esterases.html, as of December 2025), of which the CE2 and CE17 families of bacterial origin have been identified as active on β-mannans (37). Some studies of the CE2 family have reported activity on both acetylated xylan, acetylated glucomannan, and galactoglucomannan, with specificity for the 3-*O*, 4-*O*, and 6-*O* positions on mannose units (25, 38, 39). The CE17 family specifically acts on 2-*O*-acetylations on β-mannans and has been described to date only for members of the Bacillota phylum (24, 25). Here, we describe and biochemically characterize an esterase pair, a CE7 and a CExxx, from *B. cellulosilyticus* DSM 14838 involved in β-mannan metabolism. The two esterases are part of a large MUL that is specifically upregulated in response to growth on β-mannans. These esterases exhibit complementary activities and include a member of a newly identified esterase family (CExxx) that removes 2-*O*-acetylations.

## Results and discussion

### Complex β-mannan induces a mannan utilization locus that includes two esterases

*B. cellulosilyticus* DSM 14838 was tested for its ability to utilize β-mannans as the sole carbon source. In agreement with previous work, we confirmed that *B. cellulosilyticus* showed substantial growth on all substrates tested (Fig. 1b), including complex AcGGM from Norway spruce and KGM (20). To investigate the molecular basis of *B. cellulosilyticus’s* ability to process β-mannan, a proteomic study was conducted. When comparing the proteomes of *B. cellulosilyticus* on AcGGM and KGM against glucose, proteins with functions compatible with β-mannan uptake and depolymerization were identified as the top 40 most abundant in the proteomes of *B. cellulosilyticus*. Among these proteins, 14 are encoded by genes located in a MUL (Fig. 1c). The *Bc*MUL includes genes encoding three β-mannanases/mannosidase (GH26A, GH26B, and GH5_7), a membrane transporter protein (MFS), two SusC/SusD-like transporter/binding proteins, a phosphorylase belonging to the GH130_1 family, two glucanases/glucosidases (GH5_2 and GH3), a carbohydrate esterase (CE7), a transcriptional protein and two hypothetical proteins (Fig. 1d). Several detected proteins are similar to those characterized in β-mannan degrading *Bacteroides* species. This includes *Bc*GH26A, which shares 60.4 % identity to GH26A in *B. ovatus* and 61.6 % identity to GH26 in *P. dorei*, and *Bc*GH26B having 32.2 % identity to GH26B in *B. ovatus* (31, 33). A lone α-galactosidase (*Bc*GH36) outside the MUL was detected at elevated abundance when grown on β-mannans; *Bc*GH36 shares 73.5 % identity with a previously characterized GH36 in *B. ovatus* (32). In addition, a predicted esterase (BACCELL_02125), hereafter referred to as *Bc*CExxx, together with an epimerase (BACCELL_02126), were also detected to be more abundant during growth on β-mannan. *Bc*CExxx is an SGNH hydrolase-type esterase domain-containing protein not previously characterized (based on Interpro searches). The genes BACCELL_02125 and BACCELL_02126 are located in a small, separate contig in the genome of *B. cellulosilyticus* DSM 14838.

A second esterase, belonging to the CE family 7, was detected in the proteome of *B. cellulosilyticus* when growing on β-mannans. Members of the CE7 family include acetyl xylan esterase and a cephalosporin-C deacetylase, with some characterized CE7s active on both such substrates and several members with broad substrate activities (40, 41). Intriguingly, despite characterized CE7s showing activity towards a variety of acetylated structures, no members of the CE7 family have until now been reported to deacetylate β-mannans.

### Substrate testing of *B. cellulosilyticus* esterases

The activity of recombinant versions of *Bc*CE7 and *Bc*CExxx were tested on a variety of structurally characterized substrates. On AcGGM with 2-*O* and 3-*O* acetylations, both esterases were active and performed a partial deacetylation of the substrate when applied individually. A complete deacetylation of this substrate was only obtained when both enzymes were combined in the same reaction (Fig. 2a). The same feature was observed when glucomannooligosaccharides were used as a substrate (*Ri*GH26-digested KGM); indeed, both esterases were required to completely remove all acetylations (Fig. S1a). *Aloe vera* mannan, which can have multiple acetylations per mannose moiety, including the 6-*O* position (42, 43), was not completely deacetylated by the pair of esterases, although each esterase separately shows activity on this substrate (Fig. S1b). This may indicate that neither esterase has specific activity for 6-*O*-acetylations or that highly acetylated mannose moieties present more steric hindrance to substrate binding in the active site, possibly preventing deacetylation by either of these esterases.

**Fig. 2.**
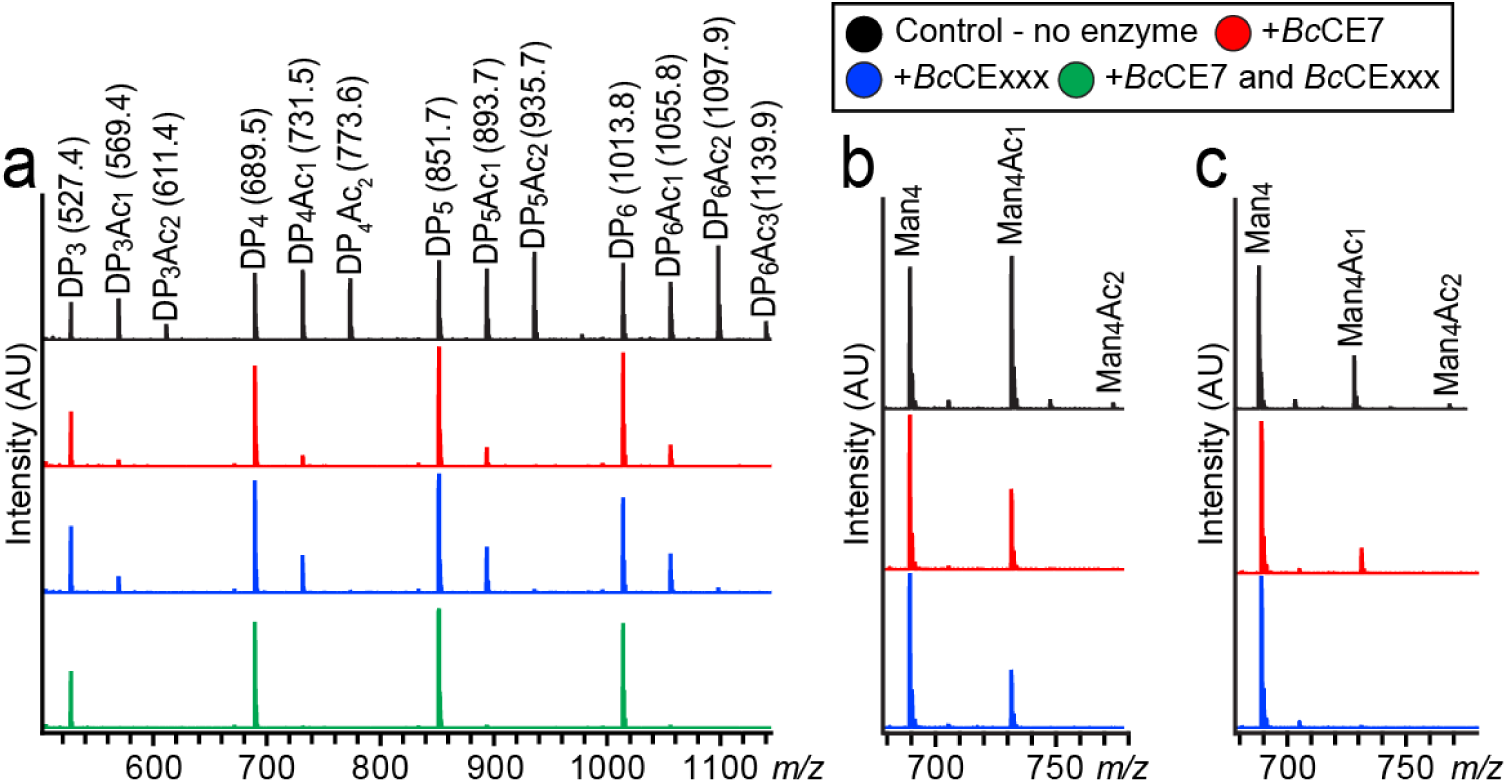
MALDI-ToF spectra of activity on β-mannan for *Bc*CE7 and *Bc*CExxx. a) AcGGM from Norway spruce was tested with the two esterases, both separately and in combination. The top row shows untreated AcGGM, while the second and third rows show treatment with *Bc*CE7 and *Bc*CExxx, respectively. Both esterases were active on the substrate, reducing peak intensities of masses corresponding to acetylated oligos. Treatments with both esterases combined (bottom row) completely removed acetylations. *Bc*CE7 and *Bc*CExxx were tested for activity on mannotetraose with acetylations in specific positions at *O*-3 and *O*-2, produced with the previously characterized *Ri*CE2 and *Ri*CE17, respectively. b) Both esterases showed some activity towards mannotetraose with 3-*O*-acetylations during the reaction time. c) *BcCExxx* and *BcCE7* act on *2-O-*acetylated mannotetraose, with *BcCExxx* completely removing all acetyl groups, while *BcCE7* shows only a minor effect. The reactions were performed with 1 µM enzyme concentration and 0.1 mg/mL substrate in 10 mM sodium phosphate pH 5.9 buffer run at 25 °C with stirring. Reactions with AcGGM and specific substrate were performed for 24 h and 1 h, respectively.

*Bc*CE7 demonstrated broad specific esterase activity, acting on several acetylated substrates, including acetylated xylan and cellulose monoacetate. It did not completely deacetylate xylan; however, a shift in the mass spectra and a higher distribution of non-acetylated xylooligosaccharides were observed (Fig. S1c). On acetylated cellulose, *Bc*CE7 completely removed all acetylations (Fig. S1d). *Bc*CExxx displayed minimal activity on acetylated xylan compared to *Bc*CE7; however, some deacetylation occurred after 24 hours (Fig. S1c). *Bc*CExxx had lower activity on acetylated cellulose, resulting in a higher ratio of mono- and di-acetylated cellooligosaccharides, but did not perform a complete deacetylation of the substrate (Fig. S1d). None of the esterases had any activity on the *N*-acetylated tetra-*N*-acetyl-chitotetraose.

### Positional specificity of the *B. cellulosilyticus* esterases

The results described above suggested that *Bc*CE7 and *Bc*CExxx may have complementary activity and are likely to be specific for different positions. To further investigate this hypothesis, we tested the enzymes’ positional preference by deacetylation of mannotetraoses with either 2-*O*- or 3-*O*-acetylations. Using transesterification reactions with characterized esterases of known specificity from *R. intestinalis* (*Ri*CE2 and *Ri*CE17) (Fig. S2) (25, 44), mannotetraoses with mainly single acetylations and, to a lesser extent, double acetylations were obtained. Both *Bc*CE7 and *Bc*CExxx were active on 3-*O*-acetylated mannotetraose with partial deacetylation during short reaction times (1 hour) (Fig. 2b). Both esterases were also active on 2-*O*-acetylated mannotetraose, whereas *Bc*CE7 displayed incomplete deacetylation, *Bc*CExxx removed all acetylations at this specific position (Fig. 2c), indicating a higher *O*-2 specificity. Interestingly, this pair of esterases in a *Bacteroides* species seems to have less positional specificity than its Bacillota species counterpart (24, 25).

Next, the ability of *Bc*CE7 and *Bc*CExxx to transacetylate mannooligosaccharides was investigated. Although reactions with *Bc*CE7 required a higher enzyme concentration (10-fold) than those with *Bc*CExxx, both esterases showed transesterification activity. On mannotetraose, *Bc*CE7 transferred mainly one acetyl group, with a small amount of two acetylations observed, while *Bc*CExxx transferred up to three acetylations (Fig. 3a and b). To investigate if *Bc*CE7 and *Bc*CExxx were capable of adding larger substituents, vinyl propionate and vinyl butyrate were also tested in transesterification reactions with mannotetraose. *Bc*CE7 transpropylated, albeit to a lesser extent than acetylate, but was unable to transbutyrylate the substrate (Fig. 3a). *Bc*CExxx transpropylated almost to the same degree as observed for acetylations, with up to three propylations per mannotetraose unit (Fig. 3b). *Bc*CExxx was also able to transbutyrylate, but mainly with one to two butyrylations (Fig. 3b). This indicates that *Bc*CExxx has more plasticity and can accommodate larger donors in transesterification reactions than previously reported for other β-mannan active esterases (25). Additionally, mannooligosaccharides from Man_2_ to Man_6_ were tested in transacetylation reactions. *Bc*CE7 failed to add acetyl groups to Man_2_, while a small amount of acetyl was added to Man_3-6_ (Fig. S3a). *Bc*CExxx transferred one acetyl to Man_2_ (with a very minor peak corresponding to two acetylations) and multiple acetyl groups to Man_3-6_ (Fig. S3b).

**Fig. 3.**
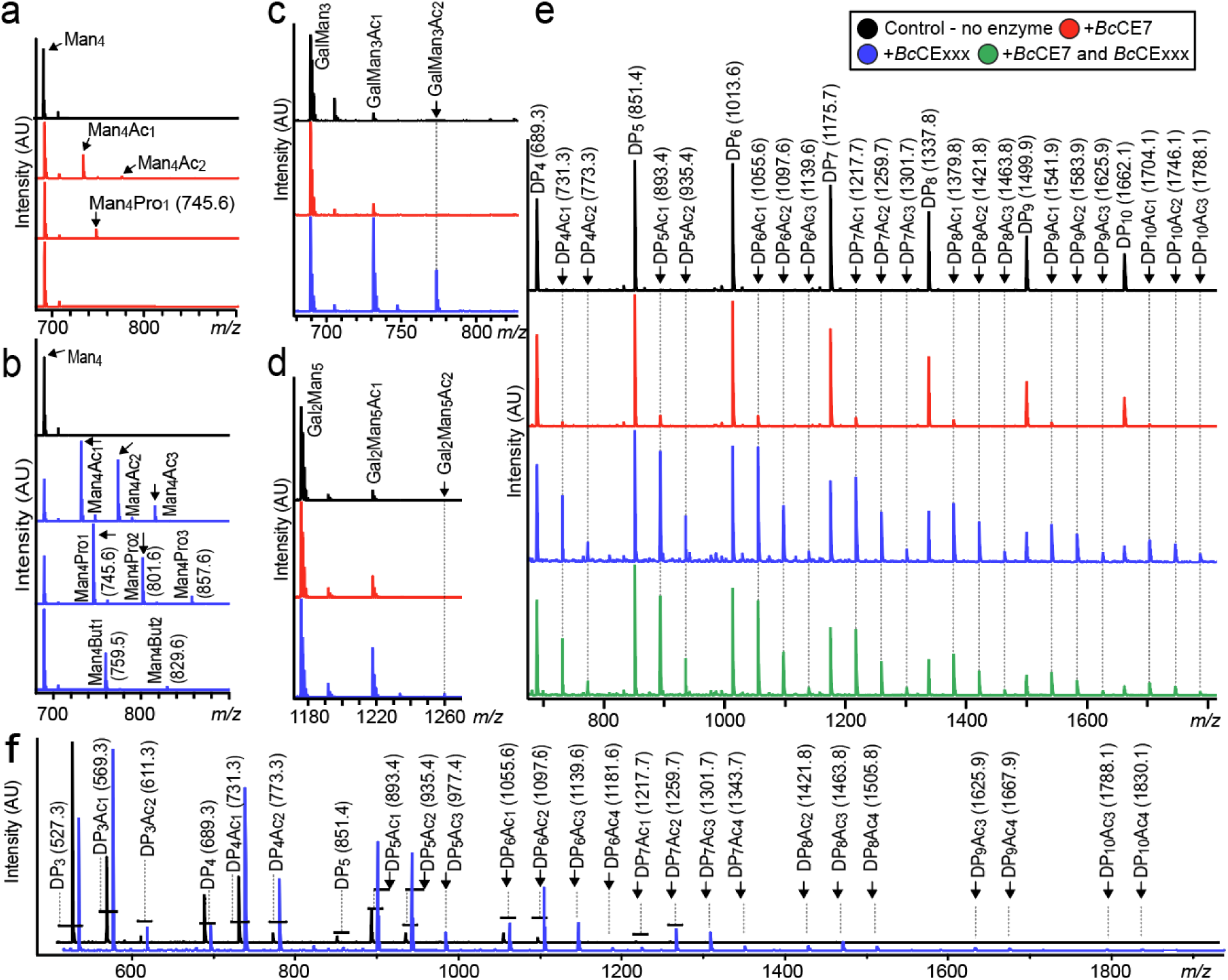
Transesterification reactions with a range of different substrates. Transesterification reactions of mannotetraose with a) *Bc*CE7 and b) *Bc*CExxx with vinyl acetate, -propionate, and -butyrate. For CE7, acetyl and propionyl (to a low extent) groups were attached to the substrate, while no transbutyrylation was observed, whereas *Bc*CExxx transesterified mannotetraose with all donors tested, although to a lesser extent for the larger donors. c) Transacetylation of GalMan_3_, d) Gal_2_Man_5_, and e) chemically deacetylated Norway spruce with *Bc*CE7 and *Bc*CExxx. f) Comparison of degradation with the endomannanase (*Ri*GH26) of KGM without (black) or with (blue) a pretreatment with *Bc*CExxx transacetylation in vinyl acetate. Transesterification reactions were conducted with 1 mg/mL substrate in 10 mM sodium phosphate (pH 5.9) and 200 nM enzyme with vinyl acetate/propionate/butyrate donors added to 50 % of the sample volume and run overnight with stirring at 25 °C. Abbreviations: Ac, acetyl; Prop, propyl; But, butyryl; Man, mannose; Gal, Galactose; *m/z*, mass/charge; DP, degree of polymerization.

Furthermore, the ability of *Bc*CE7 and *Bc*CExxx to transacetylate more complex substrates was tested by using 6^1^-α-D-galactosyl-mannotriose (GalMan_3_), 6^3^, 6^4^-α-D-galactosyl-mannopentaose (Gal_2_Man_5_), chemically deacetylated spruce mannan, polymeric KGM, and xylotriose. On GalMan_3_, *Bc*CE7 added one acetyl group, while *Bc*CExxx added two acetyl groups, but to a slightly lesser extent than on mannotriose (Fig. 3c). For Gal_2_Man_5_, both *Bc*CE7 and *Bc*CExxx added mainly one acetyl group, however, a small amount of double-acetylations was observed for *Bc*CExxx (Fig. 3d). This indicates that galactosylations reduce the activity of *Bc*CE7 and that multiple adjacent galactose substitutions prevent the full activity of *Bc*CExxx. For the non-acetylated spruce GGM, *Bc*CE7 produced a small amount of single acetylated substrates, while *Bc*CExxx added up to two acetylations on oligosaccharides with a degree of polymerization (DP) 4-5 and three acetylations on DP 6-10, demonstrating activity on longer oligosaccharides (Fig. 3e). *Bc*CExxx transacetylated polymeric KGM, which is reported to be in the range of 200-2000 kDa, indicating that its activity is not limited to oligosaccharides (14). A comparison of KGM with or without a pretreatment with *Bc*CExxx transacetylation in vinyl acetate clearly shows a profound increase in acetylations on the endomannanase *Ri*GH26 released oligosaccharides from the *Bc*CExxx transacetylated KGM (Fig. 3f).

To obtain further insight into enzyme specificity, we made comparisons with the reciprocal transacetylation reactions from *Bc*CE7 and *Bc*CExxx, followed by deacetylation with *Ri*CE2 and *Ri*CE17, which have previously been demonstrated to remove 3-*O*- and 2-*O*-acetylations, respectively (Fig. S2) (25). *Ri*CE2 also removes 4-*O*-acetylations and has some activity towards 6-*O*-acetylations. Notably, some remaining acetylation was observed after combined treatments with *Ri*CE2 and *Ri*CE17 on the *Bc*CE7 transacetylated substrate (Fig. S2c). Since there were very minor amounts of remaining acetylations of the combined treatment of *Ri*CE2 and *Ri*cE17, we may speculate that *Bc*CE7 transacetylates the reducing end mannose. *Ri*CE17, which has a strict 2-*O* specificity (25), removes almost all acetylations on transacetylated products of *Bc*CExxx (Fig. S2d). Such observations complement the findings detailed in Fig. 2.

NMR experiments were conducted to further understand which positions on the substrate *Bc*CExxx preferentially act upon. Due to the different degrees of acetylation, the ^1^H-NMR spectrum of *Bc*CExxx transacetylated mannotetraose (Fig.S4, Table S1) showed a plethora of overlapping signals compared to mannotetraose (Fig. S5, Table S2). However, three distinct new multiplets, 5.47, 5.49, and 5.53 ppm, could be identified (Fig. 4).

**Fig. 4.**
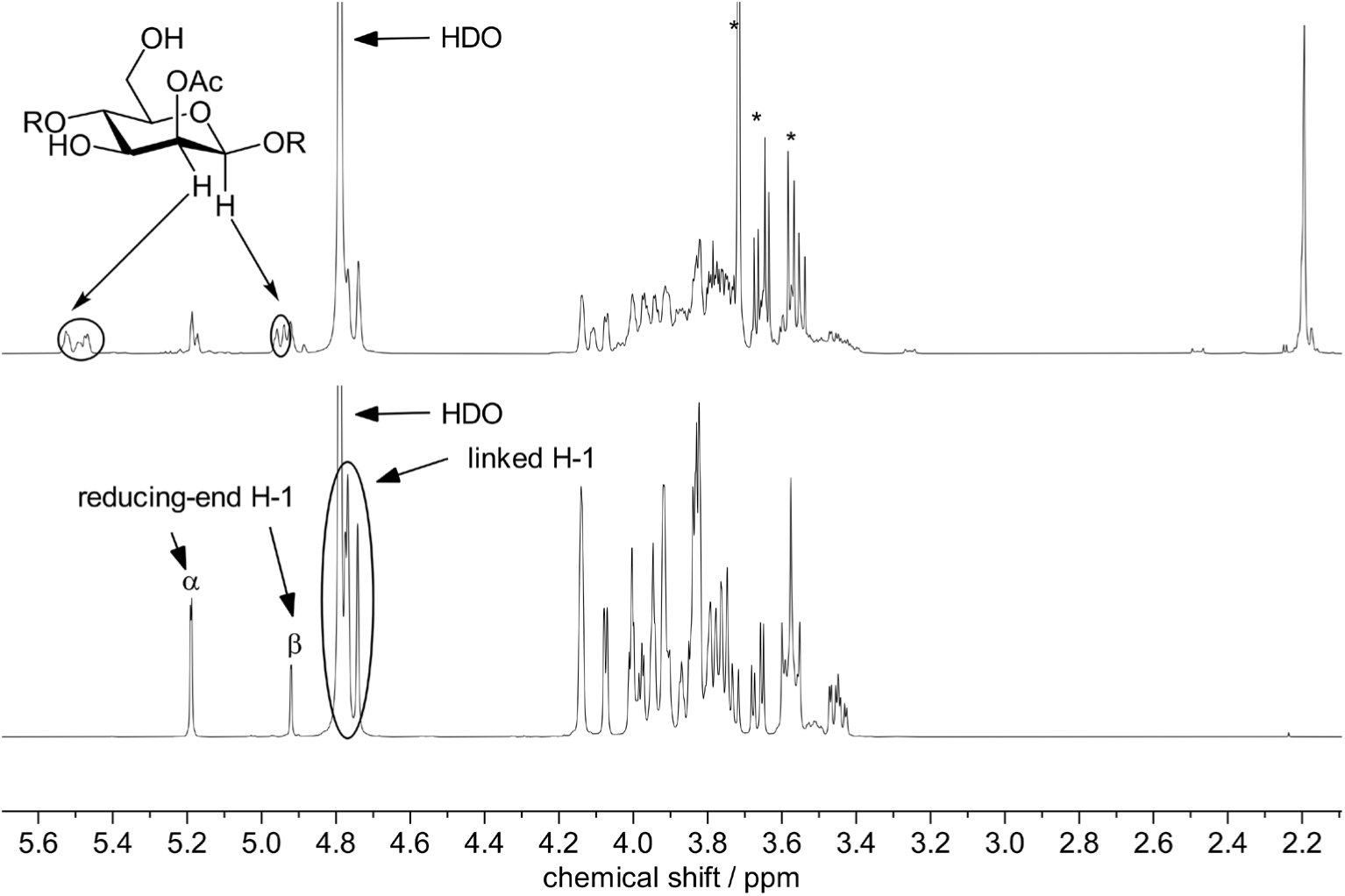
^1^H-Spectrum (400 MHz, 298 K) of mannotetraose after (top) and before (bottom) enzymatic transacetylation by *Bc*CExxx. The signal at 2.19 indicated the presence of unbound acetate. The introduction of the acetyl group at *O*-2 shifts the resonance frequencies of nearby protons downfield by ∼0.2 ppm for H-1 and ∼1.4 ppm for H-2.

We next used 2D NMR experiments to further elucidate the acetylation patterns of the products. The multiplets at 5.47 and 5.53 showed a correlation to the carbonyl peaks belonging to the acetyl groups at 176.5 ppm in HMBC (Fig. S4a). The degree of acetylation on Mannose-2 was too low to observe a correlation in HMBC. COSY spectroscopy showed that the multiplets also correlated with the anomeric protons between 4.93-4.96 ppm (Fig. S4b), confirming they belong to the protons attached to C-2 of the pyranoses, overall indicating selective acetylation at *O*-2. Additionally, the anomeric protons belonging to the acetylated products showed only correlations to carbon signals at 102.1 ppm in HSQC indicating that the reducing end 2-*O* is not acetylated since the reducing end anomeric carbons resonate at a lower frequency of 96.7 ppm (Fig. S4c, d, e). Furthermore, the multiplet at 5.45-5.48 ppm showed a correlation to the carbon signal at 69.9 ppm in HSQC-TOCSY (Fig. 5) and the proton signal at 3.60 ppm in ^1^H-TOCSY (Fig. S4f). Both signals belong to C-4 and H-4 of the non-reducing end mannose, indicating that acetylation at *O*-2 of the non-reducing end mannose did indeed occur. The chemical shifts for the protons indicating 2-*O*-acetylation were in agreement with previously reported values (25). A full assignment of proton and carbon signals for mannotetraose and its transacetylation product is given in the supporting information (Table S1 and S2). Acetyl migration was observed during the acquisition of NMR-spectra of the transacetylated products (45). While ^1^H- and HSQC-NMR spectra acquired immediately after dissolving the sample in D_2_O only showed 2-*O*-acetylation, spectra acquired one day later showed signals indicating 3-*O*-acetylation and 4-*O*-acetylation (Fig. S4j and k). The chemical shifts of these signals agreed with previously reported data for acetylated mannotriose (25). The predicted structures of both esterases were further investigated. The AlphaFold structures of both esterases from the AlphaFold Protein Structure Database have an average confidence measure (pLDDT), without signal peptides, of 97.21 and 97.46 for *Bc*CE7 and *Bc*CExxx, respectively (46, 47). *Bc*CE7 was superimposed with the crystal structure of two characterized CE7s from *Bacillus subtilis* (PDB: 1ODS, RMSD = 1.27 Å) and *Paenibacillus* sp. R4 (PDB: 6AGQ, RMSD = 1.27 Å), which have been shown to have activity on acetylated xylan or xylooligosaccharides (Fig. S6a) (40, 41). The *Bc*CE7 has a highly similar structure to the other two CE7s, including the catalytic triad Ser281-Asp363-His392, with an additional unique N-terminal fold that has no annotations (based on Interpro searches) (Fig. S6b). We next compared the structure of *Bc*CExxx with that of *Ri*CE17 (PDB: 6HFZ), the only known structure of an esterase with similar activity towards 2-*O*-acetylations (25), resulting in an RMSD of 2.61 Å. While the overall 3D structures of the two esterases are different (Fig. S6c), they have a spatially similar catalytic triad (Ser37-His241-Asp238 in *Bc*CExxx and Ser41-His193-Asp190 in *Ri*CE17) and an Asn (Asn134 in *Bc*CExxx and Asn110 in *Ri*CE17) forming an oxyanion pocket (Fig. S6d). Contrary to *Ri*CE17, which has a CBM35 in its structure that forms a clamp crucial for substrate binding and catalytic activity, *Bc*CExxx has a narrower loop containing a tryptophan (Trp88) in close proximity to the tryptophan in the CBM35 in *Ri*CE17 (Trp326), potentially aiding substrate stacking. The higher turnover rate and transesterification of larger substituents could be explained by a more flexible catalytic site and the narrower clamp formed by the tryptophan-containing loop.

**Fig. 5.**
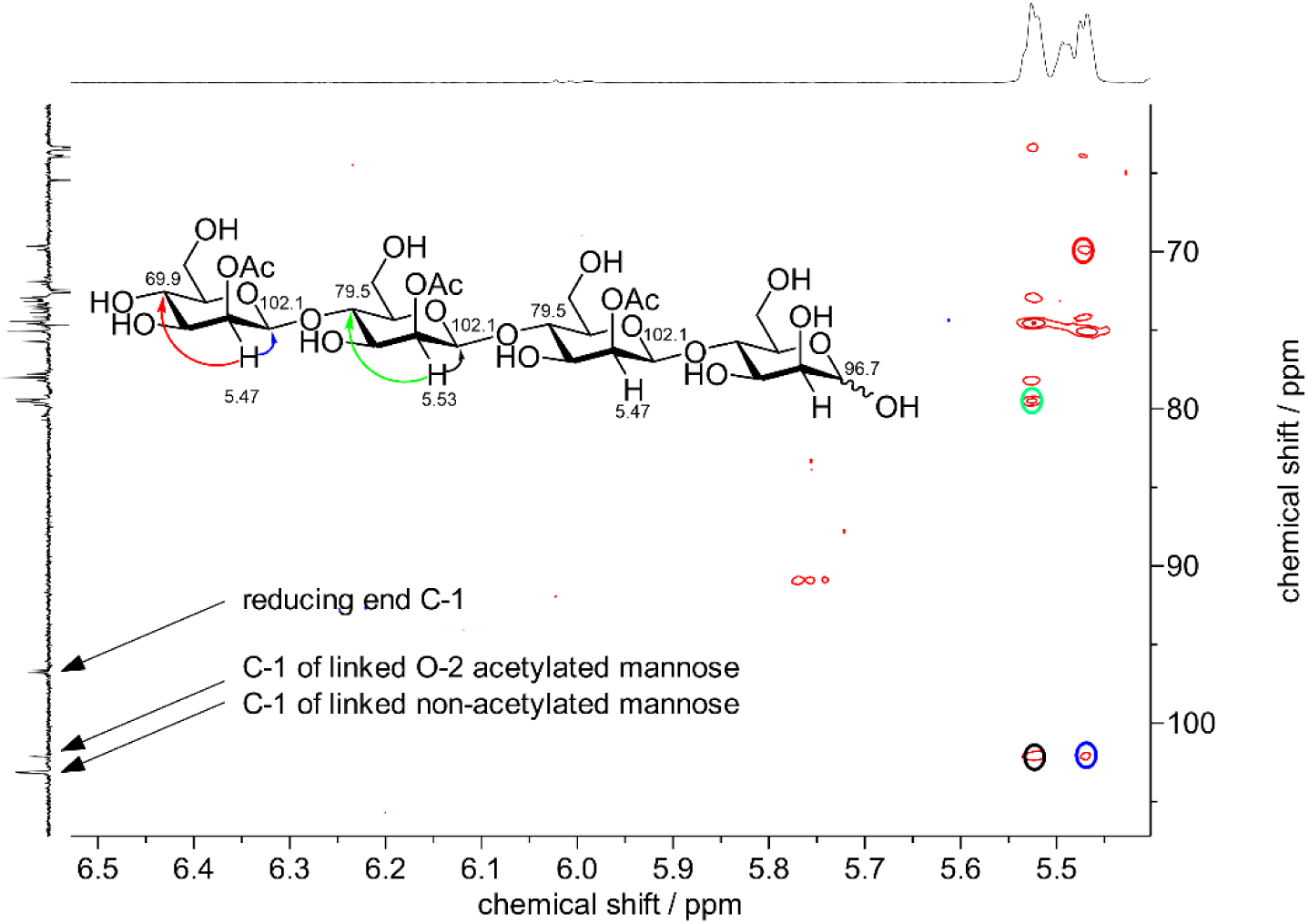
HSQC-TOCSY spectrum (D_2_O, 400 MHz, 298 K) showing acetylation of the non-reducing-end mannose. Horizontal axis: ^1^H-NMR (full spectrum in Fig. S4g.) Vertical axis: DEPTQ (Full Spectrum in Fig. S4h). The non-reducing-end mannose can be easily differentiated from the in-chain mannoses by the lower chemical shift of C-4 (69.9 ppm vs. 79.5 ppm, represented by red and green circles/arrows, respectively). Furthermore, anomerically linked mannoses can be differentiated from the reducing-end mannose by the chemical shift of their respective anomeric carbons (C-1). C-1 of non-acetylated mannose resonated at 103.1 ppm, C-1 of *O*-2 acetylated mannose resonated at 102.1 ppm (based on the correlations represented by black and blue circles/arrows, respectively), and the reducing end C-1 resonates at a much lower chemical shift of 96.8 ppm. The full HSQC-TOCSY spectrum is given in Fig. S4i.

### Catalytic turnover and pH optimum

The pH and temperature optimum were analyzed with pNP-acetate, a widely used substrate in esterase activity assays. For both enzymes, there were small differences in the range of pH 6.5-7.5 (Fig. S7), and they were active from 20 to 50 °C. *Bc*CE7 had high activity in the range of pH 6.75 to 7.25, although at pH 8 the initial acetate release was higher, the total release in prolonged reactions was lower. A similar feature for *Bc*CE7 was also observed at the highest temperatures (45-50 °C). *Bc*CExxx showed the highest activity at pH 7.25, and both enzymes displayed high activity at 30-37 °C.

The turnover rate was determined using *Ri*GH26-digested spruce AcGGM at pH 7.25 and 37 °C. The activity of *Bc*CExxx on spruce mannan was 10-fold greater than that of *Bc*CE7 (Table 1). Compared with *Ri*CE2 and *Ri*CE17 (Table S3), *Bc*CExxx has a 5-fold faster turnover rate, while *Bc*CE7 shows the slowest deacetylation rate. Furthermore, it should be noted that *Bc*CExxx has a markedly lower activity on pNP-acetate, which is almost 1/200 compared to AcGGM. This implies that, in general, for enzymes such as these, pNP-acetate is not a very suitable substrate for studying overall activity. On an additional note, *Bc*CE7 demonstrated a 17-fold higher turnover rate on pNP-acetate compared to *Bc*CExxx. This finding shares a similar relationship as previously observed between CE2 and CE17 in *R. intestinalis* (25), further suggesting that pNP-acetate is a more suitable substrate for testing the activities of enzymes active on equatorial rather than axially oriented acetyl groups.

**Table 1.** Turnover rates of the esterases of *B. cellulosilyticus* on spruce AcGGM and pNP-acetate. The AcGGM was pre-digested with the *Ri*GH26 endomannanase. The turnover number was determined based on triplicate measurements of the amount of acetate released linearly within the first hour. For the deacetylation of pNP-acetate, a total of six replicates from two independent experiments were used for the determination of the turnover number. An enzyme concentration of 50 nM was used for the AcGGM reactions, while 50 nM of *Bc*CE7 and 100 nM of *Bc*CExxx were used in the pNP-acetate reactions. All reactions were run in phosphate buffer at pH 7.25 and 37 °C.

| Deacetylation of Norway spruce AcGGM | Kcat[s<sup>-1</sup>] |
| --- | --- |
| *BcCE7* | 20.75 (± 1.802) |
| *BcCExxx* | 201.2 (± 27.21) |
| *BcCE7* + *BcCExxx* | 107.3 (± 4.195) |
| Deacetylation of pNP-acetate |  |
| *BcCE7* | 20.11 (± 3.46) |
| *BcCExxx* | 1.161 (± 0.252) |

Herein, we have characterized an esterase pair from a commensal gut *Bacteroides* that performs cooperative removal of *O*-acetylations. The metabolic commitment of two predominant phyla in our guts, Bacillota and Bacteroidota, to produce functionally mannan active esterases suggests that acetylation of β-mannans may be more prevalent in our diets than commonly recognized. This study adds significant contributions by providing data on catalytic rates on natural substrates. Notably, the *Bc*CExxx has a profoundly higher catalytic turnover rate than previously characterized 2-*O*-acetyl specific esterases. Transacetylation reactions indicate that *Bc*CExxx is active on both oligosaccharides and polymeric substrates, acetylating all mannose residues except the reducing end. Based on the results presented here, as well as previous studies on β-mannan degradation, we propose a β-mannan degradation pathway of a gut *Bacteroides* representative (Fig. S8).

## Materials and Methods

### Substrates

All carbohydrate stocks were prepared at 10 mg/mL in double-distilled water (ddH_2_O) or in specified buffer and filter sterilized using a 0.22-µm membrane filter (Sarstedt AG & Co., Germany).

#### (i) Polysaccharides

Konjac glucomannan was purchased from Megazyme International (Wicklow, Ireland). *A. vera* mannan (Acemannan polysaccharide) was purchased from Elicityl (Crolles, France). Cellulose monoacetate (degree of acetylation of 0.6) was a kind gift from Qi Zhou, KTH Royal Institute of Technology, Stockholm.

#### (ii) Oligo- and monosaccharides

Glucose (G1) was purchased from Sigma-Aldrich (St. Louis, MO, USA), while mannobiose (Man_2_), mannotriose (Man_3_), mannotetraose (Man_4_), mannopentaose (Man_5_), mannohexaose (Man_6_), xylotriose, and tetraacetyl-chitotetraose were purchased from Megazyme. Hydrolyzed versions of konjac glucomannan and *A. vera* mannan were generated in-house using the endo-mannanase *Ri*GH26 (23) in 10mM sodium phosphate (pH 5.9). Reactions were incubated for 16 h at 37 °C following removal of the enzyme using a Vivaspin 20 filtration unit (10,000-molecular-weight-cutoff [MWCO] polyethersulfone [PES]; Sartorius). Carbohydrates were subjected to lyophilization on an ALPHA 2-4 LD Plus freeze dryer (Christ, Germany) and stored as solids. Acetylated galactoglucomannan (AcGGM) from Norway spruce and acetylated xylan from birch were produced in-house from dried wood chips as described previously (22, 48). The AcGGM was treated with Multifect ® Xylanase (Genencor-Danisco) and nano-filtrated to diafilter the released xylose residues.

#### (iii) Transesterification of substrates

Oligosaccharides were dissolved in 10 mM sodium phosphate (pH 5.9) to 1 mg/mL and enzymes were added to a final concentration of 200 nM (2 µM for *Bc*CE7). Vinyl acetate/propionate/butyrate (Sigma) was added to 50% of the sample volume. The samples were incubated at 25 °C with 600 rpm stirring overnight in a thermomixer (Eppendorf, Norway). Samples were frozen at -20 °C, resulting in the vinyl acetate/propionate/butyrate liquid remaining on top of the frozen aqueous phase, which was then removed. Samples were thawed on ice and immediately filtered through a pre-washed 1 mL Amicon Ultracel 3kDa ultrafiltration device (Merck KGaA, Germany) to remove the enzymes and avoid any de-esterification reactions. For transacetylated polymeric KGM, absolute ethanol was added (to 80% final concentration) to the frozen aqueous phase to quench the esterase activity.

The ethanol was evaporated, and the dry pellet was resuspended in buffer. The substrate was further hydrolyzed with 1 µM *Ri*GH26 at 25 °C with stirring. For NMR, a scaled-up reaction was conducted to produce 5 mg of transacetylated mannotetraose. 5 mL of 1 mg/mL mannotetraose was used in a 50 mL Falcon tube, following the same protocol as described. The sample was incubated with shaking at 250 rpm overnight at 25 °C and filtered with a pre-washed Vivaspin 20 filtration unit (10,000-molecular-weight-cutoff [MWCO] polyethersulfone [PES]; Sartorius) before freeze-drying. Transacetylated versions of mannotetraose with *Ri*CE2 and *Ri*CE17 were made as described previously (25).

### Bacterial strains and culture conditions

*B. cellulosyliticus* DSM 14838 (49) was routinely cultured in freshly prepared minimal medium (MM) supplemented with 5 mg/mL glucose (50). All fermentations were carried out at 37 °C in an anaerobic cabinet (Whitley A95 Workstation; Don Whitley, UK) under an atmosphere of 85% N2, 5% CO2, and 10% H2. Growth was assessed by measuring the optical density at 600 nm [OD600] at regular intervals for up to 24 h. All growth experiments were performed in triplicate.

### Analysis of the *B. cellulosilyticus* proteome

Triplicate cultures of *B. cellulosilyticus* were grown in minimal medium (50) supplemented with either 0.5% (w/v) glucose or β-mannan (AcGGM or KGM) as the sole carbon source. Samples (10 mL) were obtained at the mid-exponential growth phase, and the cell pellets were collected by centrifugation at 4500 × g for 10 min at 4 °C following resuspension in lysis buffer (50 mM Tris-HCl pH 7.5, 200 mM NaCl, 0.1% v/v Triton x-100). Cell lysates were prepared using a bead-beating approach, whereby glass beads (diameter ≤ 106 μm) were added to the samples and cells disrupted by 3 × 60 s cycles of bead-beating, using FastPrep24 instrument (MP Biomedicals, Santa Ana, CA, USA). Cell debris was removed by centrifugation at 16600 × g for 20 min and proteins were precipitated overnight in 16% ice-cold TCA. Finally, proteins were dissolved in 30 μL 50mM Tris-HCl, pH 8.4. For each sample, 25 μL of proteins were processed using an S-Trap™ 96-well plate digestion protocol (Protifi, Fairport NY, USA), according to the manufacturer’s instructions. The resulting peptides were analyzed on a nanoLC-MS/MS system, consisting of a nanoElute UHPLC connected to a Tims-ToF Pro ion-mobility mass spectrometer (both from Bruker), equipped with a nano-electrospray ion source. Peptides were separated using a PepSep Reprosil C18 reverse-phase (1.5 µm, 100Å) 25 cm X 75 μm analytical column coupled to a ZDV Sprayer (Bruker Daltonics, Bremen, Germany). The column was maintained at 50 °C using the integrative oven. Equilibration of the column was performed before the samples were loaded (equilibration pressure 800 bar). The flow rate was set to 300 nL/min and the samples were separated using a solvent gradient from 5% to 25% solvent B over 70 min, and to 37% over 9 min. The solvent composition was then increased to 95% solvent B (0.1% [v/v] formic acid in acetonitrile) over 10 min and maintained at that level for an additional 10 min, for a total separation run time of 99 min. Solvent A consisted of 0.1% [v/v] formic acid in milliQ water. The timsTOF Pro was run in positive ion data-dependent acquisition PASEF mode with the control software Compass Hystar version 5.1.8.1 and timsControl version 1.1.19. The acquisition mass range was set to 100 – 1700 *m/z*.

Mass spectrometry (MS) raw data were analyzed with FragPipe v19.0 and searched against the *B. cellulosilyticus* DSM 14838 protein sequence database (GCA_000158035.1, 5837 protein entries) with MSFragger (51). The database was supplemented with common contaminants, such as human keratin, trypsin, and bovine serum albumin, in addition to reversed sequences of all protein entries for estimation of false discovery rates (FDR) with Philosopher (52). Carbomidomethylation of cysteine residues was used as fixed modification, while oxidation of methionine and protein N-terminal acetylation were used as variable modifications. Trypsin was selected as proteolytic enzyme, one maximum missed cleavage site was allowed and matching tolerance levels for both MS and MS/MS were 20 ppm. The results were filtered to achieve a protein 1% FDR and quantification was done using IonQuant including normalization between samples and the feature ‘match between runs’ to maximize protein identifications (53). Perseus v1.6.2.3 was used for further analysis (54). A protein was considered valid if it was detected in at least two of the three biological replicates in at least one glycan substrate. Proteins identified as potential contaminants were removed. Missing values were imputed from a normal distribution (width of 0.3 and downshifted 1.8 standard deviations from the original distribution) in total matrix mode and differential abundance analysis was performed using an unpaired two-tailed Student’s t-test with a permutation-based FDR set to 0.05. Heat maps were generated using Perseus v1.6.2.3.

### Plasmid design, CE7 and CExxx overexpression and purification

The plasmid constructs to produce the recombinant *Bc*CE7 (EEF89986.1) and *Bc*CExxx (EEF90239.1) were generated by GenScript Biotech through a synthesized DNA fragment introduced into pET28a(+) and pET30a(+), respectively. The gene sequences were designed to exclude the N-terminal signal peptide (predicted by the SignalP v5.0 server (55)) and added a histidine tag for purification through immobilized metal affinity chromatography (IMAC). Constructs were transformed into chemically competent Escherichia coli BL21 STAR cells (Invitrogen), and precultures were inoculated to 1% in 500 mL tryptone yeast extract (TYG) containing 50 mg/mL kanamycin, followed by incubation of the culture for 16 h at 23 °C. Protein overexpression was induced by adding isopropyl β-D-thiogalactopyranoside (IPTG) to a final concentration of 200 mM. Recombinant *Bc*CE7 and *Bc*CExxx production continued overnight at 23 °C, after which the cells were collected by centrifugation. Cells were collected by centrifugation at 8,000 x *g* for 10 min at 4 °C, and pellets were stored at −80 °C until further processing. Proteins were purified using standard IMAC purification procedures, employing a Vibracell ultrasonic homogenizer (Sonics and Materials, USA) to lyse cells (24). Eluted protein fractions were pooled, concentrated using a Vivaspin 20 centrifugal concentrator (10-kDa molecular weight cutoff), and applied to a HiLoad 16/600 Superdex 75 pg gel filtration column (GE Healthcare). Pure protein samples were buffer exchanged to remove imidazole against 10mM Tris-HCl (pH 7.0) and concentrated as described above. Protein purity was determined by sodium dodecyl sulfate-polyacrylamide gel electrophoresis (SDS) analysis. Protein concentration was determined by measuring A280 absorbance and converting to molarity using calculated extinction coefficients.

### Activity assays

Unless otherwise stated, enzyme reactions were carried out in 10 mM sodium phosphate (pH 5.9) and 0.1 mg/mL substrate. The activities of *Bc*CE7 and *Bc*CExxx were tested at concentrations from 1 to 10 μM. Reaction mixtures were preheated (25 °C for 10 min) in a Thermomixer C incubator with a heated lid (Eppendorf), before the addition of the enzyme (in a final volume of 100 µL) for further incubation (up to 24 h) at 25 °C and 700 rpm. Reaction products were then analyzed by matrix-assisted laser desorption ionization-time of flight mass spectrometry (MALDI-ToF MS) on an Ultraflex MALDI-ToF/ToF MS instrument (Bruker Daltonics, Germany) equipped with a 337-nm-wavelength nitrogen laser and operated by the MALDI FlexControl software (Bruker Daltonics). Samples were prepared by combining 1 μL of reaction mixture with 2 μL matrix (0.9% 2,5-dihydroxybenzoic acid (DHB)–30% acetonitrile [vol/vol]) directly applied on an MTP 384 target plate (Bruker Daltonics, Germany), and dried under a stream of warm air. All measurements were performed in positive ion, reflector mode with 1,000 shots taken per spectrum.

### NMR

^1^H-NMR and ^13^C-NMR spectra were recorded at 298 K on a Bruker AVANCE III HD 400 MHz instrument equipped with BBFO room temperature probe. Chemical shifts are reported in ppm with Na-Trimethylsilylpropanoate-*d*_4_ (δ ^1^H/^13^C = 0.00 ppm) as external standard. Multiplicities are reported as follows: s = singlet, d = doublet, t = triplet, q = quartet, m = multiplet. Coupling constants (J) are reported in Hz. The following pulse programs where used: “zg30” for 1H, “deptqgpsp.2” for DEPTQ-13C, “cosyqf45” for 1H-1H COSY, “dipsi2gpphzs” for TOCSY “hsqcetgpsi2” for HSQC, “shqqcetgpsisp2.2” for 2D-selective HSQC, “selhsqcgpsisp” for 1D-selective HSQC, “hsqcdietgpsisp.2” for HSQC-TOCSY and “hmbcetgpl3nd” for HMBC. 5 mg of *Bc*CExxx transacetylated mannotetraose was produced (as described above), which was then lyophilized, redissolved in D_2_O (1 mL) to deuterate exchangeable protons and lyophilized again. The freeze-dried sample was then dissolved in D_2_O and a 1H-spectrum and an HSQC-spectrum were acquired as fast as possible to mitigate any migration of acetyl groups (25, 56).

### Structure analysis

The predicted AlphaFold structures of *Bc*CE7 and *Bc*CExxx were obtained from the AlphaFold Protein Structure Database (47). The average confidence measure (pLDDT) was calculated with signal peptides removed and is 97.21 for *Bc*CE7 and 97.46 for *Bc*CExxx. Superimposition of structures and figures was made using PyMOL version 3.0.3.

### pH and temperature optimum

For the determination of pH and temperature optima, the enzymes were assayed in 100 mM sodium phosphate buffer with 1 mM pNP-acetate. For the pH optimum, reactions were carried out at 37 °C in buffers ranging from pH 6.0 to 8.0. To determine the optimal temperature, reactions were carried out at pH 7.25 with temperature varying from 21.8 °C (room temperature) to 50 °C. Final enzyme concentrations of 50 nM *Bc*CE7 and 100 nM of *Bc*CExxx were employed. The assays were initiated by adding 10 µL of enzyme or buffer solution with a multichannel pipette to 90 µL of assay solution within a pre-tempered 96-well plate in triplicate, and the absorbance was measured at 350 nm for 15 min in 15 to 20 second intervals in a plate reader. The absorbance of p-nitrophenol at 350 nm is almost unaffected by its protonation state. Standard curves between 0 and 1 mM of p-nitrophenol were measured under the same pH and temperature conditions. For data analysis, the individual absorbance measurements were normalized by subtracting the mean of the buffer controls, converted to concentrations based on the respective standard and the initial slope was fitted targeting a coefficient of determination of 0.99. The concentration changes were used to calculate the k_cat_.

### Quantification of acetate release

To determine the acetate release rates on a natural substrate, a solution of 100 mg/mL *Ri*GH26-digested AcGGM in 100 mM sodium phosphate pH 7.25 was prepared. Reactions were initiated in triplicate by adding the esterases either alone to a final concentration of 50 nM or in a mixture of 25 nM each and incubated at 37 °C and 700 rpm shaking on a thermomixer. Samples of 50 µL were withdrawn immediately after enzyme addition and after 5, 10, 20, 30, 40, 50, and 60 min, mixed with 50 µL 1:1 acetonitrile and isopropanol and heated to 100 °C for 2 min. After centrifuging and filtering the samples, the acetate content was analyzed on a REZEX ROA-Organic Acid H_+_ 300×7.8mm ion exclusion column (Phenomenex, USA) attached to an RSLC Ultimate 3000 HPLC (Dionex, USA). 10 µL of sample was injected, eluted by an isocratic flow (0.6 mL/min) of 5 mM H_2_SO_4_ mobile phase and detected with a UV detector at 210 nm.

The degree of acetylation of the substrate was checked by treatment in the presence of 100 mM NaOH at 4 °C for one hour. The acetate standard was prepared between 0 and 80 mM of acetic acid in the substrate solution. Acetylation control and standard were mixed with acetonitrile and isopropanol and processed the same way as the samples. For data analysis, the area of the acetate peak was measured, converted to acetate concentrations based on the standard and the initial slope was fitted targeting a coefficient of determination of 0.99. k_cat_ values were derived from the concentration changes.

### Statistics and reproducibility

All experiments were carried out in biological duplicates or triplicates.

## Acknowledgments

This study acknowledge financing from the research council of Norway (RCN); project number 244259. Lars Jordhøy Lindstad was supported by a Faculty funded PhD project and had additional funding from RCN 309558. Essential lab facilities to conduct this project was financed by the RCN Infrastructure grants 270038, 295910, 296083

## Notes

### Competing Interest Statement

The authors have declared no competing interest.

