## Supplementary file for "Deacetylation of β-Mannans by Two Complementary Carbohydrate Esterases from the Human Gut Microbe *Bacteroides cellulosilyticus*"

Supplemental information

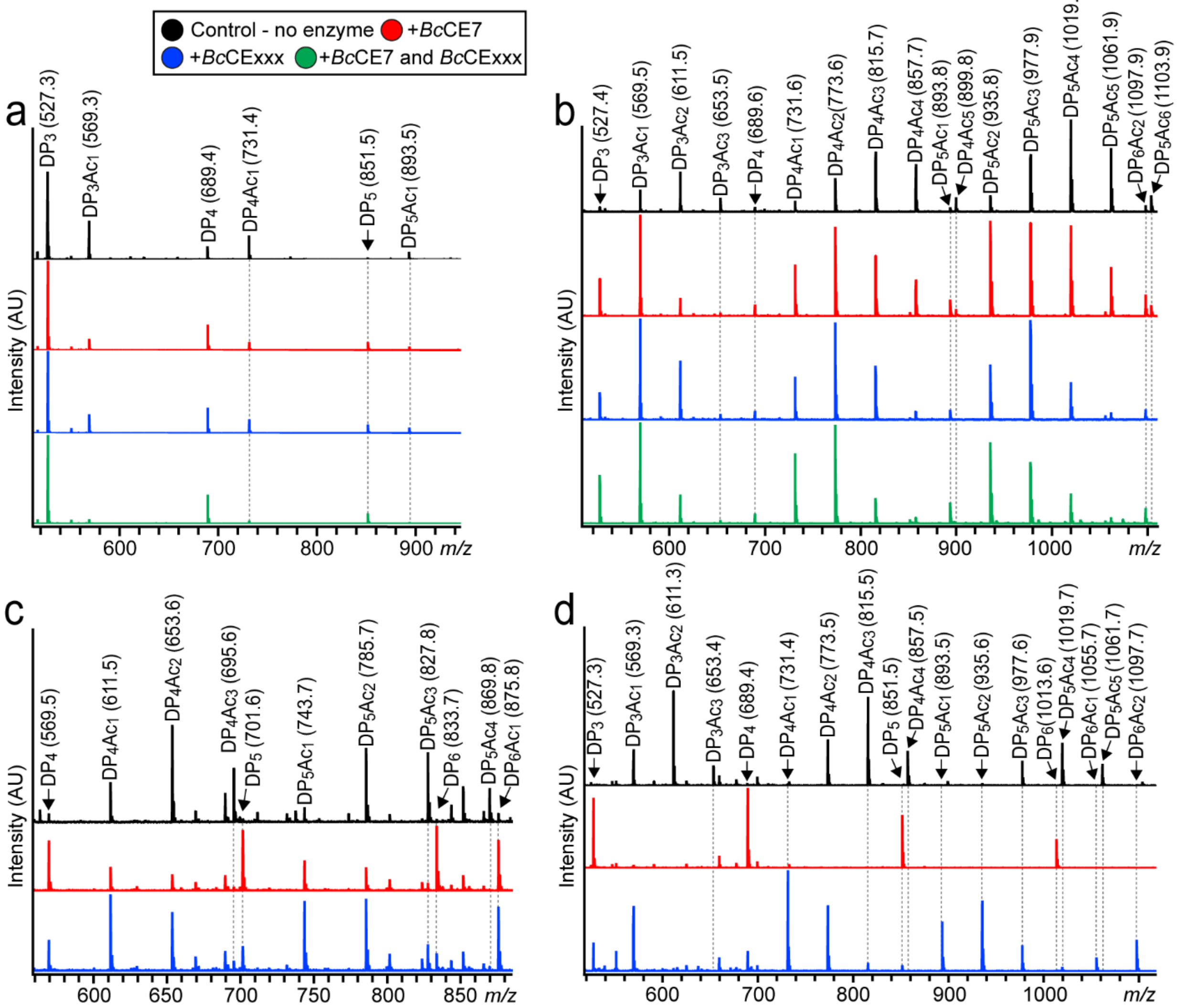


Fig. S1. MALDI-ToF spectra of enzyme reactions by *Bc*CE7 and *Bc*CExxx on various substrates. Deacetylation of *Ri*GH26-digested KGM (a) and *A. vera* mannan (b). Deacetylation of acetylated xylan (c) and cellulose monoacetate (d). The reactions were carried out with 1 µM enzyme concentration and 0.1 mg/mL substrate in 10 mM sodium phosphate pH 5.9 buffer at 25 °C with stirring for 24 h. Abbreviations: Ac, acetyl; DP, degree of polymerization; m/z, mass/charge.

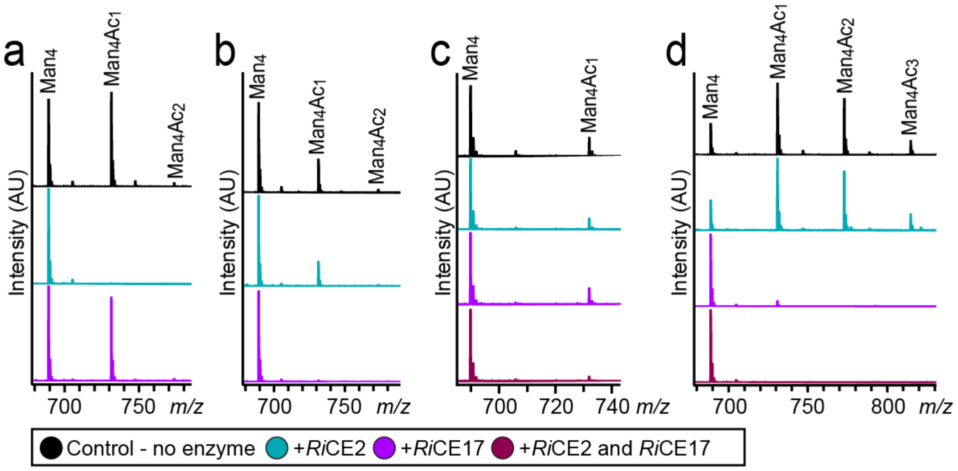


Fig. S2. Test of *Ri*CE2 and *Ri*CE17 in deacetylation reactions on transacetylated products; Deacetylation of mannotetraose transacetylated with *Ri*CE2 (a) and *Ri*CE17 (b) (to control that no acetyl migration occurred during the production of these substrates), and with *Bc*CE7 (c) and *Bc*CExxx (d). The reactions were carried out with 1 µM enzyme concentration and 0.1 mg/mL substrate in 10 mM sodium phosphate pH 5.9 buffer at 25 °C with stirring for 1 h. Abbreviations: Ac, acetyl; Man, mannose; m/z, mass/charge.

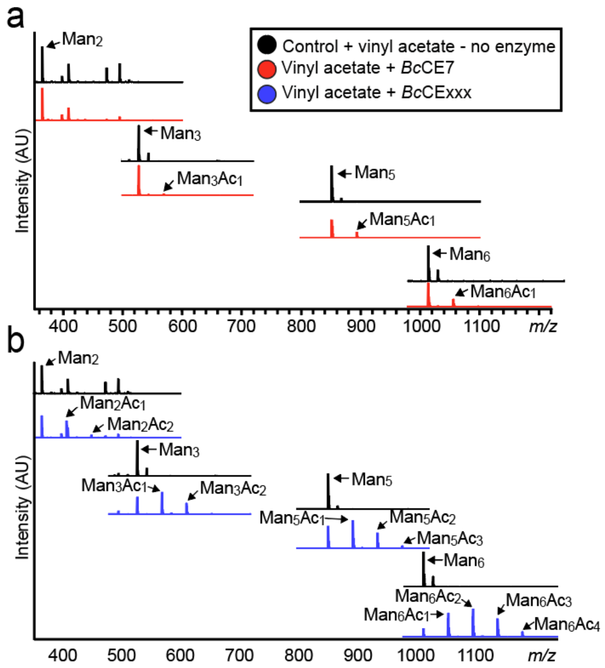


Fig. S3. a) Transacetylation of Man$_2$, Man$_3$, Man$_5$, and Man$_6$ with *Bc*CE7. No acetyl group was observed for Man$_2$, while mainly one acetylation was added to the larger mannooligosaccharides. b) *Bc*CExxx transacetylated all tested substrates, with multiple acetylations for each. Transesterification reactions were conducted with 1 mg/mL substrate in 10 mM sodium phosphate (pH 5.9) and 200 nM enzyme with vinyl acetate donors added to 50 % of the sample volume and run overnight with stirring at 25 °C. Abbreviations: Ac, acetyl; Man, mannose; *m/z*, mass/charge.

#### NMR Data of acetylated mannotetraose products

Table S1. Peak assignments for acetylated mannotetraose units.

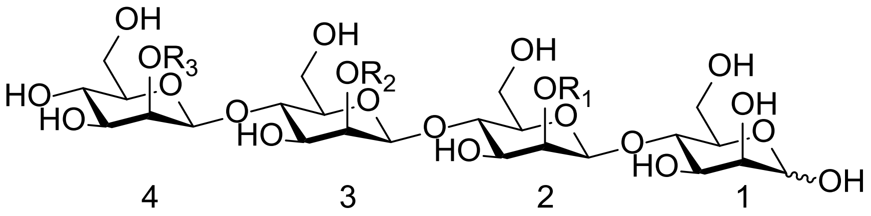


| | Mannose-unit | | |
|---|---|---|---|
| | 4<br>$R_3$ = Ac | 3<br>$R_2$ = Ac | 2<br>$R_1$ = Ac |
| 1 H | 4.94 | 4.96 | 4.92 |
| C | 102.1 | 102.1 | 102.1 |
| 2 H | 5.47 | 5.53 | 5.49 |
| C | 75.1 | 74.5 | 74.6 |
| 3 H | 3.86 | 4.02 | 3.97 |
| C | 74.2 | 73.0 | 73.4 |
| 4 H | 3.60 | 3.87 | 3.85 |
| C | 69.9 | 79.5 | 79.5 |
| 5 H | 3.53 | 3.64 | 3.54 |
| C | 78.0 | 78.2 | 77.9 |
| 6 $H_1$ | 3.72 | 3.78 | 3.78 |
| $H_2$ | 3.95 | 3.95 | 3.95 |
| C | 64.0 | 63.5 | 63.5 |

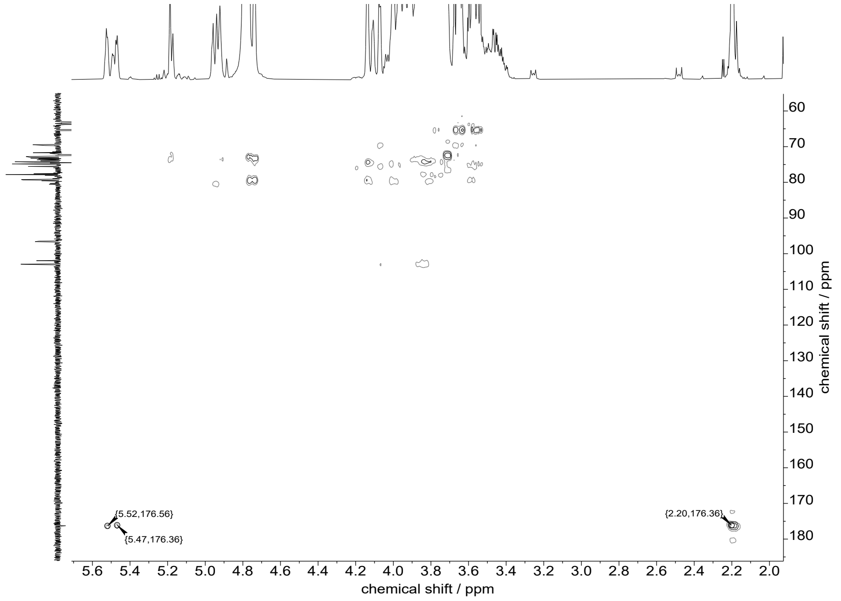


Fig. S4 a. HMBC spectrum of acetylated mannotetraose products in $D_2O$.

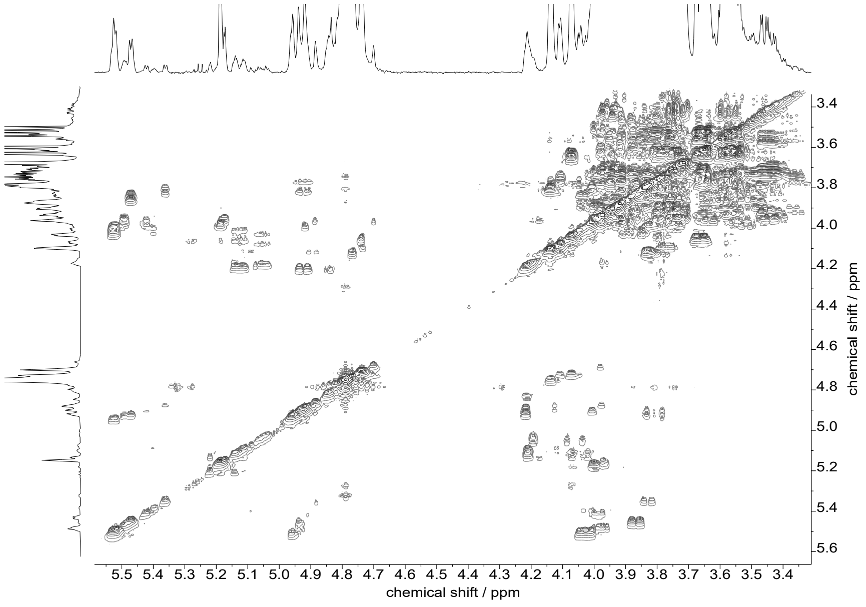


Fig. S4 b. $^1$H-$^1$H COSY spectrum of acetylated mannotetraose products in $D_2O$.

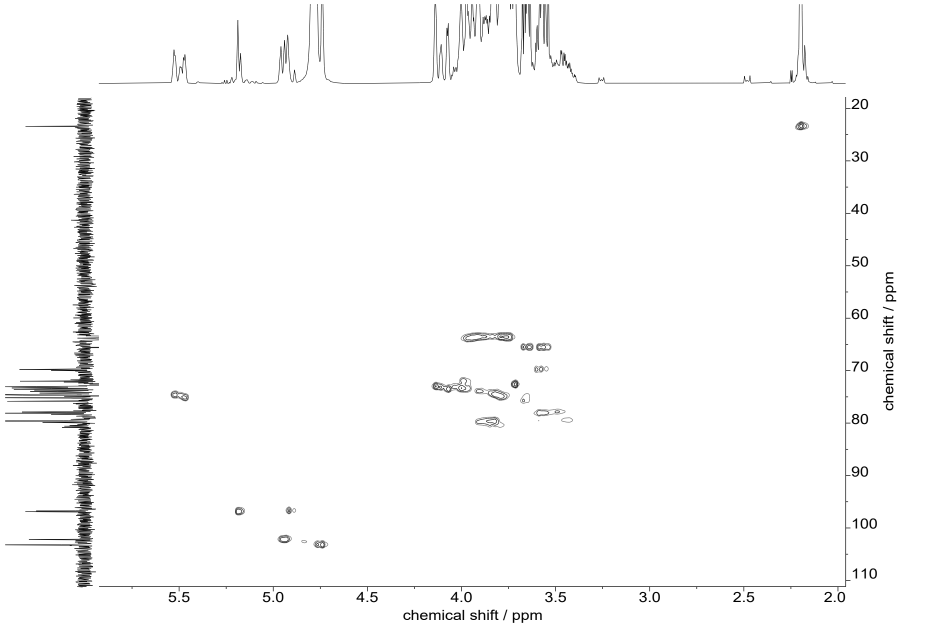


Fig. S4 c. HSQC spectrum of acetylated mannotetraose products in $D_2O$.

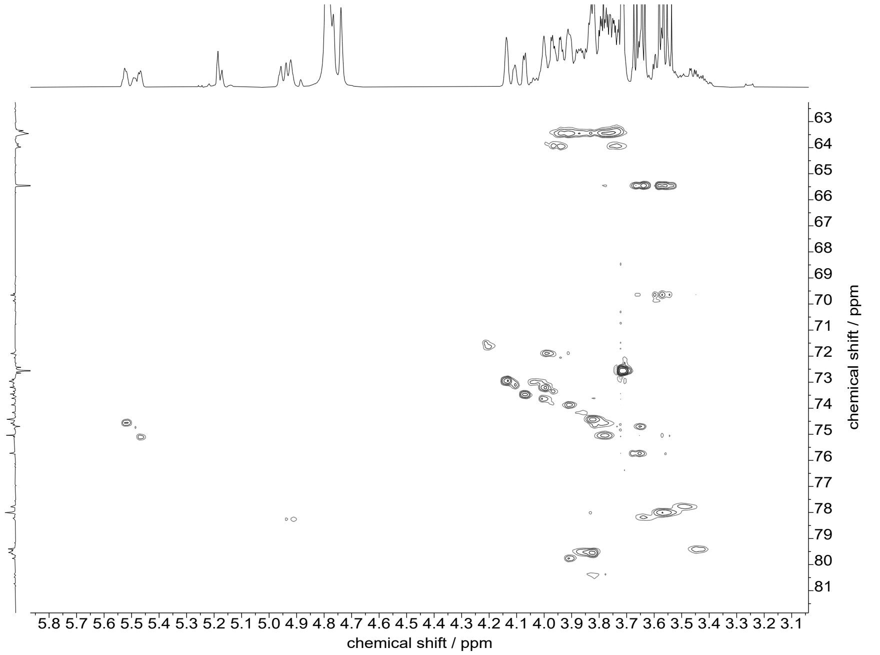


Fig. S4 d. 2D-selective HSQC spectrum of acetylated mannotetraose products in $D_2O$.

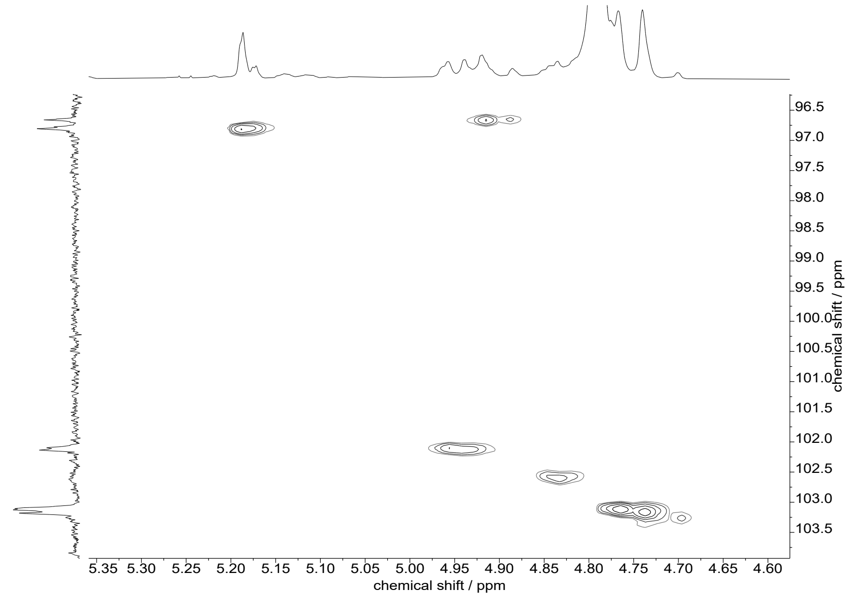


Fig. S4 e. 2D-selective HSQC spectrum of acetylated mannotetraose products in $D_2O$.

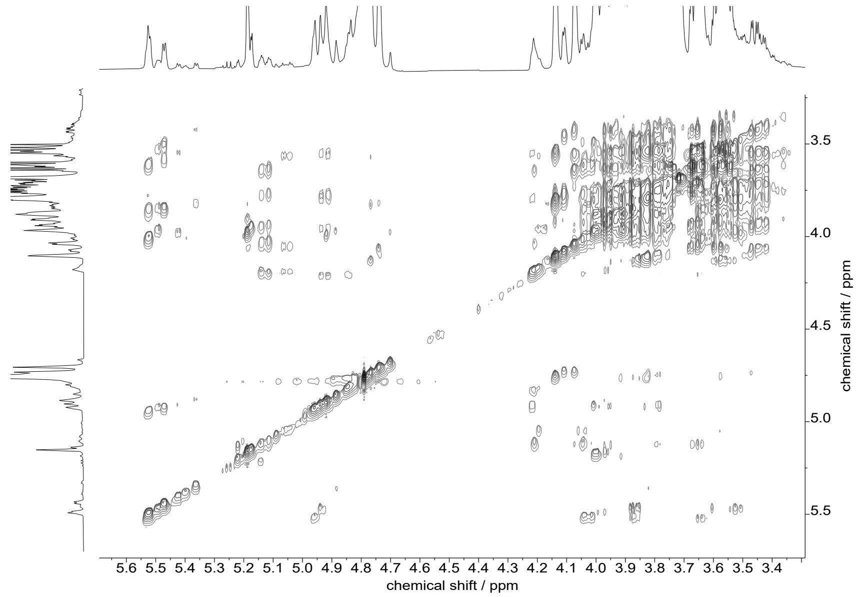


Fig. S4 f. TOCSY spectrum of acetylated mannotetraose products in $D_2O$.

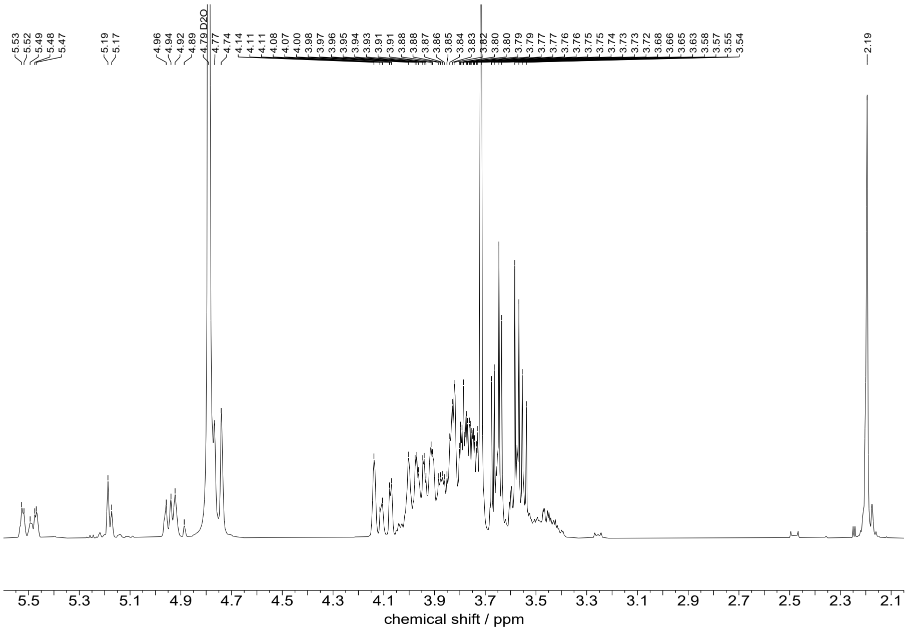


Fig. S4 g. [1]H-NMR spectrum (400 MHz, $D_2O$) of acetylated mannotetraose products.

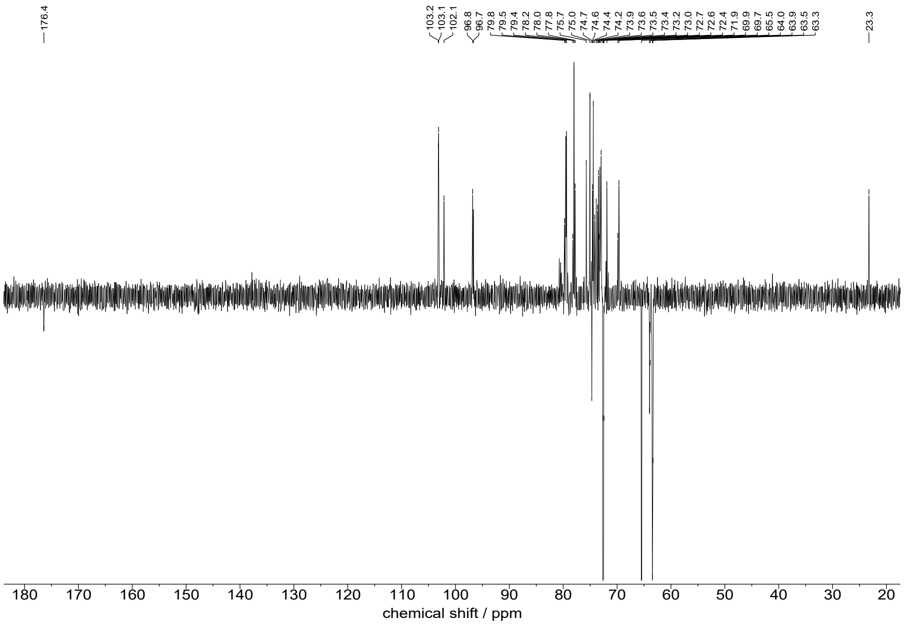


Fig. S4 h. DEPTQ spectrum (100 MHz, $D_2O$) of acetylated mannotetraose products.

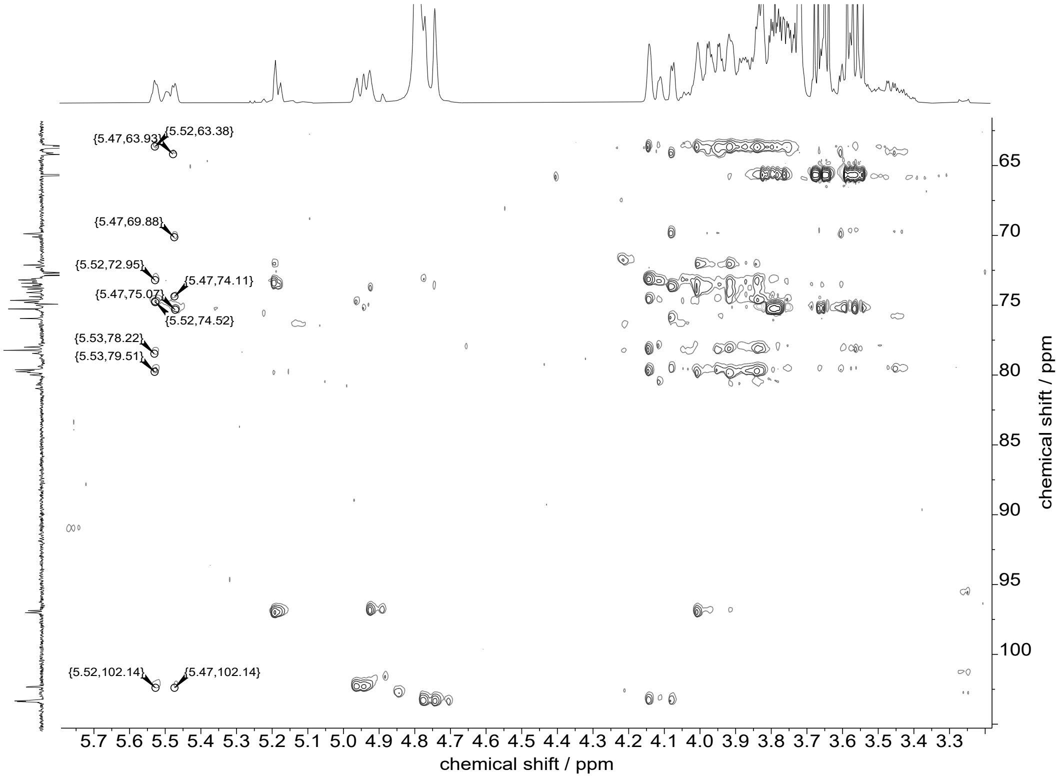


Fig. S4 i. HSQC-TOCSY spectrum of acetylated mannotetraose products in $D_2O$ (full spectrum of the partial spectrum shown in Fig 4).

### Acetyl Migration

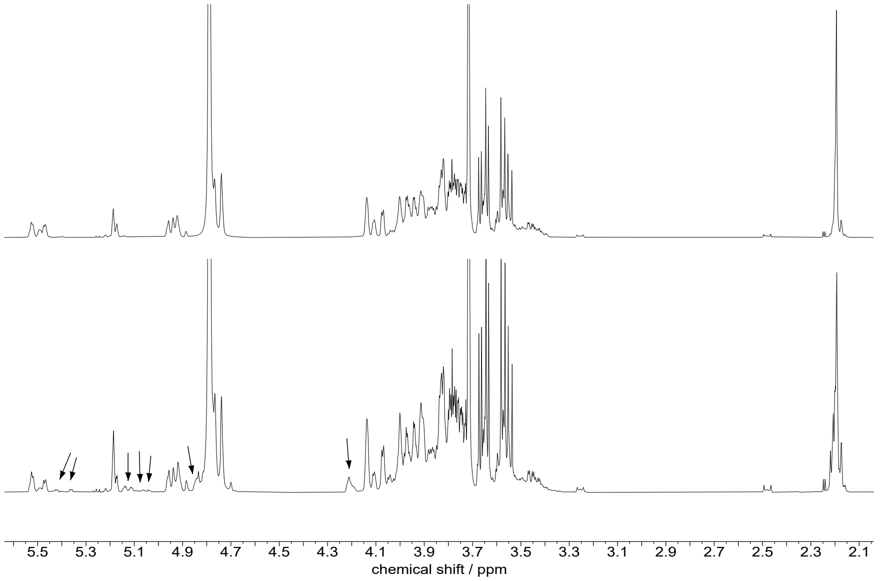

Fig S4 j. $^1$H-NMR spectrum (400 MHz, $D_2O$) of transacetylated mannotetraose before (upper spectrum) and after (lower spectrum) acetyl migration. New signals are marked by arrows.

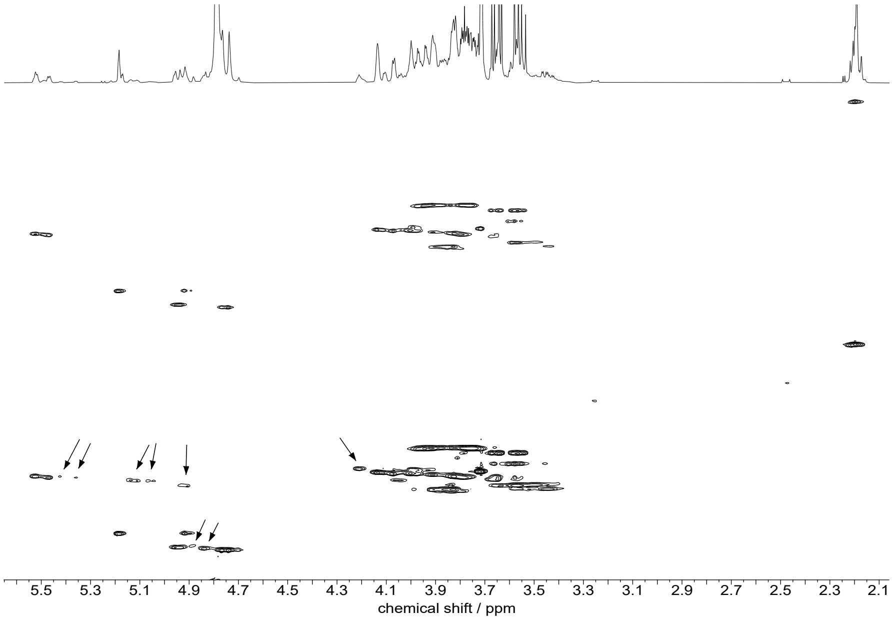


Fig S4 k. HSQC spectrum before (top) and after (bottom) acetyl migration in $D_2O$. New signals are marked by arrows.

#### NMR Data of Mannotetraose

Table S2. Peak assigments for mannotetraose.

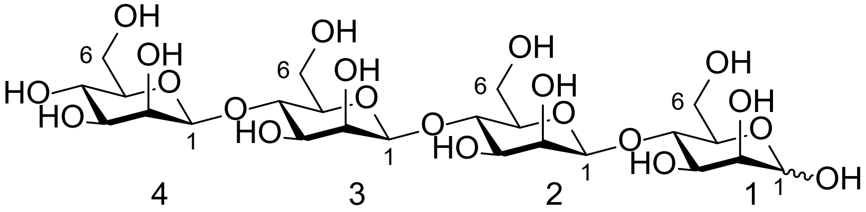

| | Mannose-unit | | | | |
|---|---|---|---|---|---|
| | 4 | 3 | 2 | 1-β | 1-α |
| 1 H<br>C | 4.77, s<br>103.2 | 4.74, s<br>103.1 | 4.74, s<br>103.1 | 4.92, d, $J^3$ = 0.7 Hz<br>96.7 | 5.19, d, $J^3$ = 1.3 Hz<br>96.8 |
| 2 H<br>C | 4.07, d, $J^3$ = 3.2 Hz<br>73.5 | 4.14<br>73.0 | 4.14<br>72.9 | 4.01<br>73.6 | 3.99<br>73.2 |
| 3 H<br>C | 3.66, dd, $J^3$ = 9.6, 3.2 Hz<br>75.7 | 3.84<br>74.4 | 3.84<br>74.4 | 3.84<br>74.6 | 3.92<br>73.8 |
| 4 H | 3.57 | 3.85 | 3.85 | 3.85 | 3.91 |
| C | 69.7 | 79.5 (4 peaks) | | | 79.7 |
| 5 H<br>C | 3.45, ddd, $J^3$ = 3.2 Hz 9.5<br>6.8 2.2 Hz<br>79.4 | 3.58<br>78.0 | 3.58<br>78.0 | 3.51<br>77.8 | 3.58<br>71.9 |
| 6 $H_1$<br>$H_2$<br>C | 3.74<br>3.96<br>64.0 | 3.79<br>3.92<br>63.5 | 3.79<br>3.92<br>63.5 | 3.79<br>3.92<br>63.5 | 3.78<br>3.86<br>63.5 |

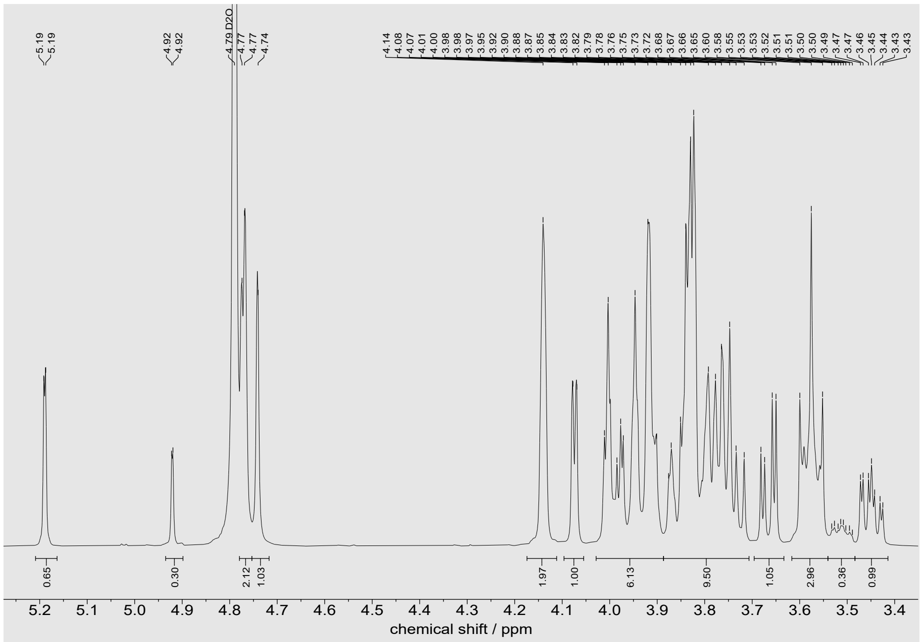


Fig. S5 a. $^1$H-NMR spectrum (400 MHz, $D_2O$) of mannotetraose.

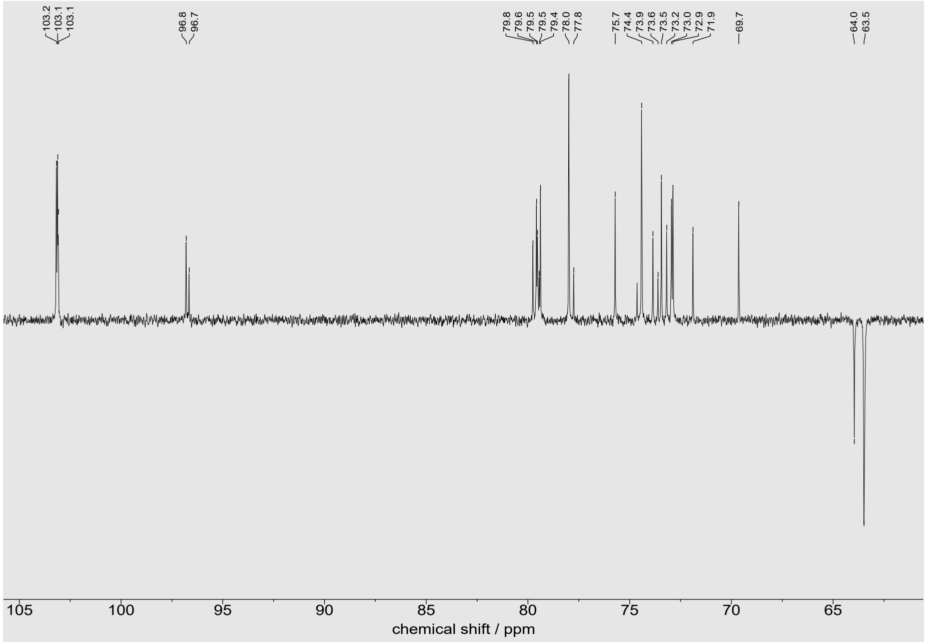


Fig. S5 b. DEPTQ spectrum (100 MHz, $D_2O$) of mannotetraose.

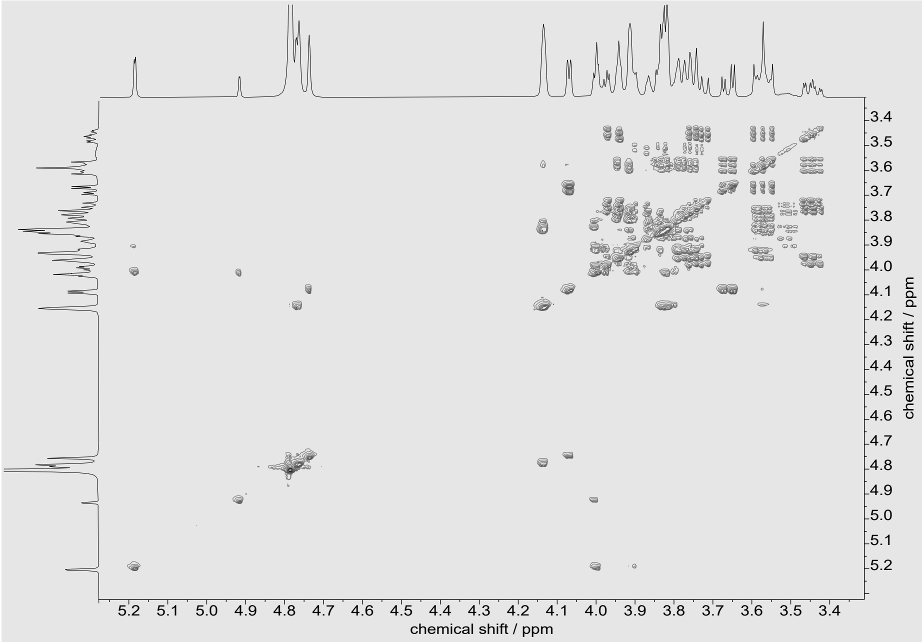


Fig. S5 c. $^1H$-$^1H$ COSY spectrum of mannotetraose in $D_2O$.

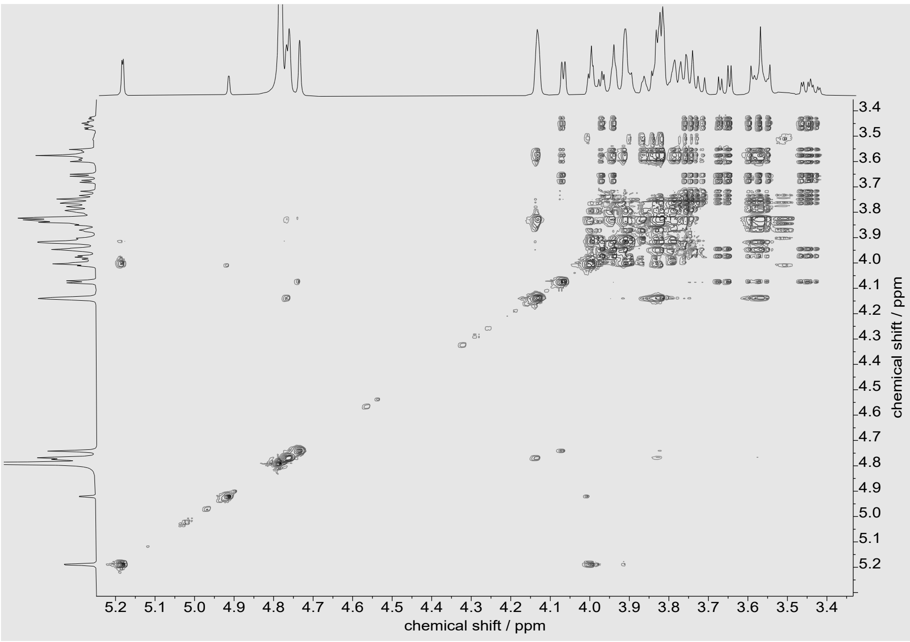


Fig. S5 d. TOCSY spectrum of mannotetraose in $D_2O$.

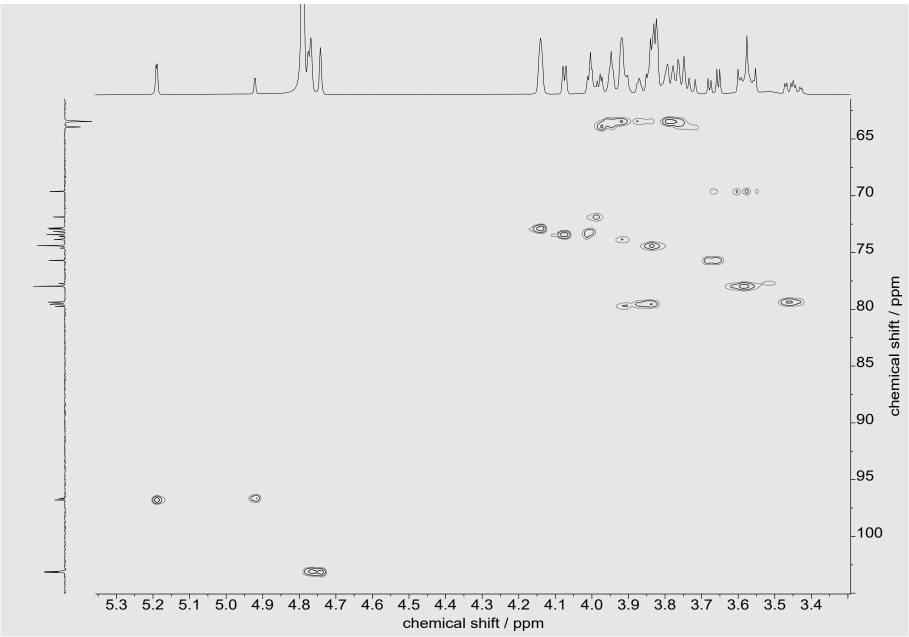


Fig. S5 e. HSQC spectrum of mannotetraose in $D_2O$.

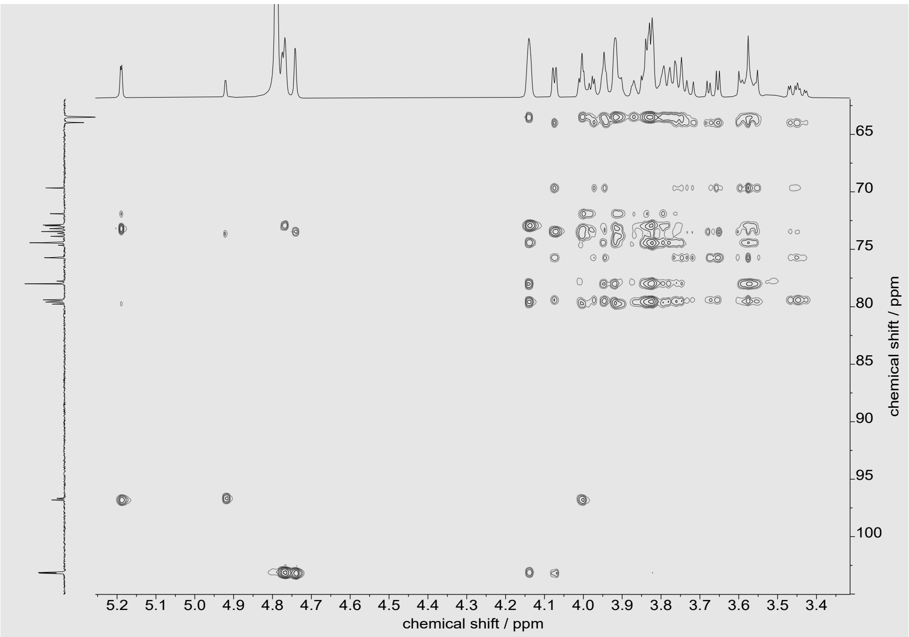


Fig. S5 f. HSQC-TOCSY spectrum of mannotetraose in $D_2O$.

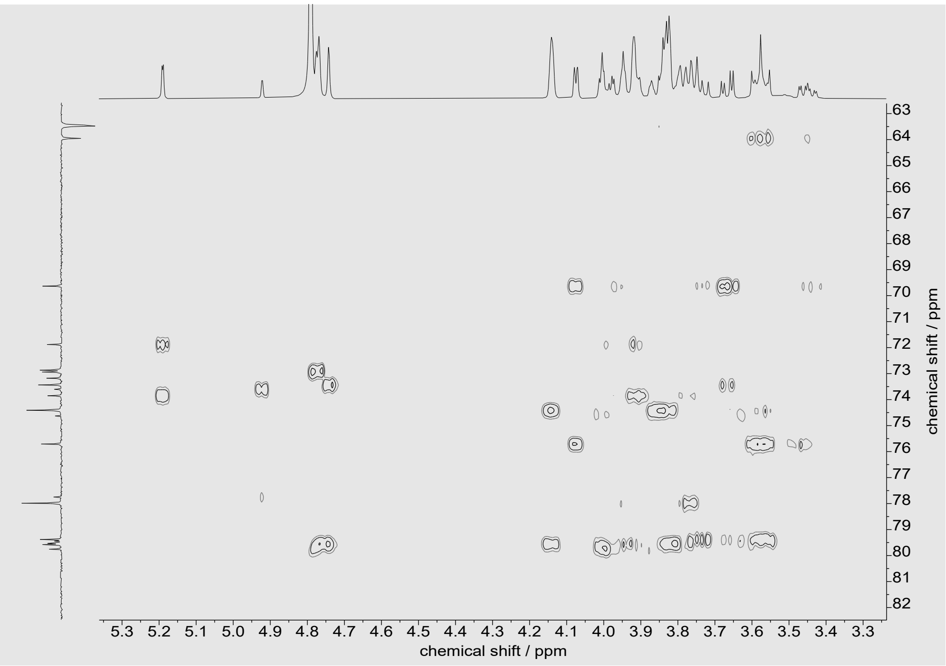


Fig. S5 g. HMBC spectrum of mannotetraose in $D_2O$.

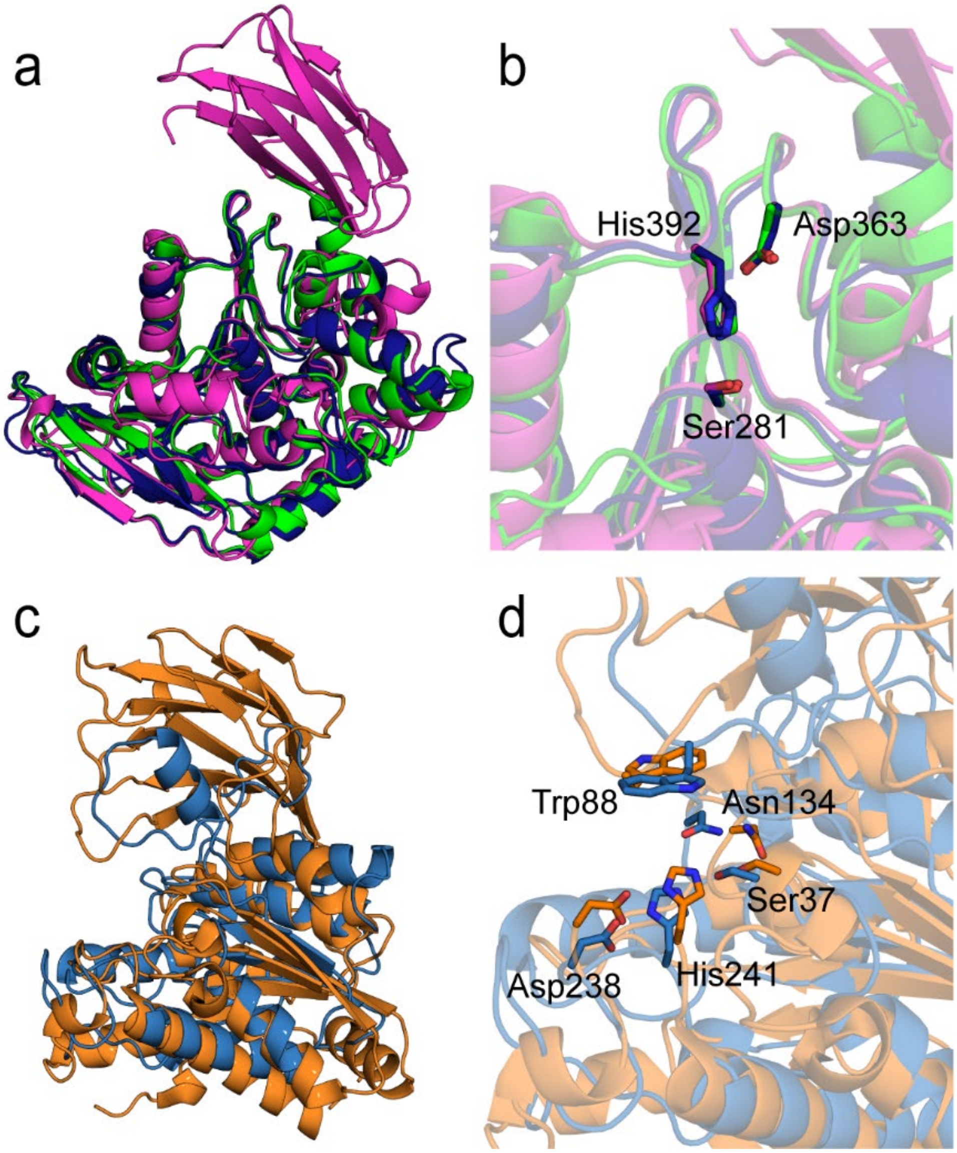


Fig. S6. Predicted AlphaFold structures of *Bc*CE7 and *Bc*CExxx. a) *Bc*CE7 (magenta) superimposed with CE7's from *Bacillus subtilis* (green) (PDB: 1ODS, RMSD = 1.27 Å) and *Paenibacillus* sp. R4 (dark blue) (PDB: 6AGQ, RMSD = 1.27 Å). b) The active site of all three CE7s with the catalytic triad Ser-His-Asp (*Bc*CE7 is numbered). c) *Bc*CExxx (blue) superimposed with *Ri*CE17 (orange) (PDB: 6HFZ) with an RMSD of 2.61 Å. d) Active site of *Bc*CExxx and *Ri*CE17. The predicted Ser-His-Asp catalytic triad of *Bc*CExxx (numbered), including the Trp for substrate stacking and Asn in the oxyanion hole, with the corresponding amino acids in *Ri*CE17 (Ser41-His193-Asp190, Trp326, and Asn110, respectively).

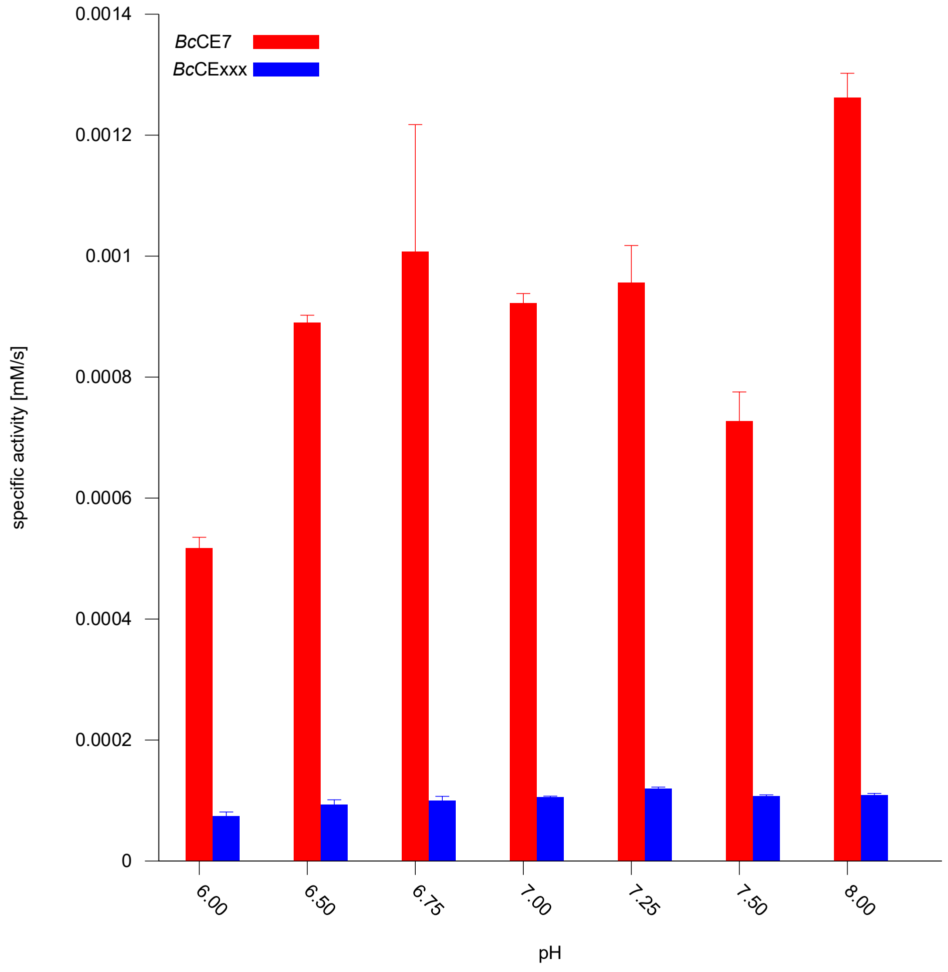


Fig. S7. pNP-acetate hydrolysis by *Bc*CE7 and *Bc*CExxx in different pH. The reactions were performed with 50 nM *Bc*CE7 and 100 nM *Bc*CExxx in 100 mM sodium phosphate buffer with 1 mM pNP-acetate at 37 °C.

Table S3. Turnover rate by *Ri*CE2 and *Ri*CE17 on *Ri*GH26-digested spruce AcGGM. The turnover number was determined based on triplicate measurements of the amount of acetate released linearly within the first hour. All reactions were run with 50 nM enzyme concentration in phosphate buffer at pH 7.25 and 37 °C.

| Deacetylation of Norway spruce AcGGM | Kcat[s-1] |
|---|---|
| *Ri*CE2 | 42.46 (± 2.89) |
| *Ri*CE17 | 38.31 (± 3.62) |
| *Ri*CE2 + *Ri*CE17 | 51.86 (± 2.47) |

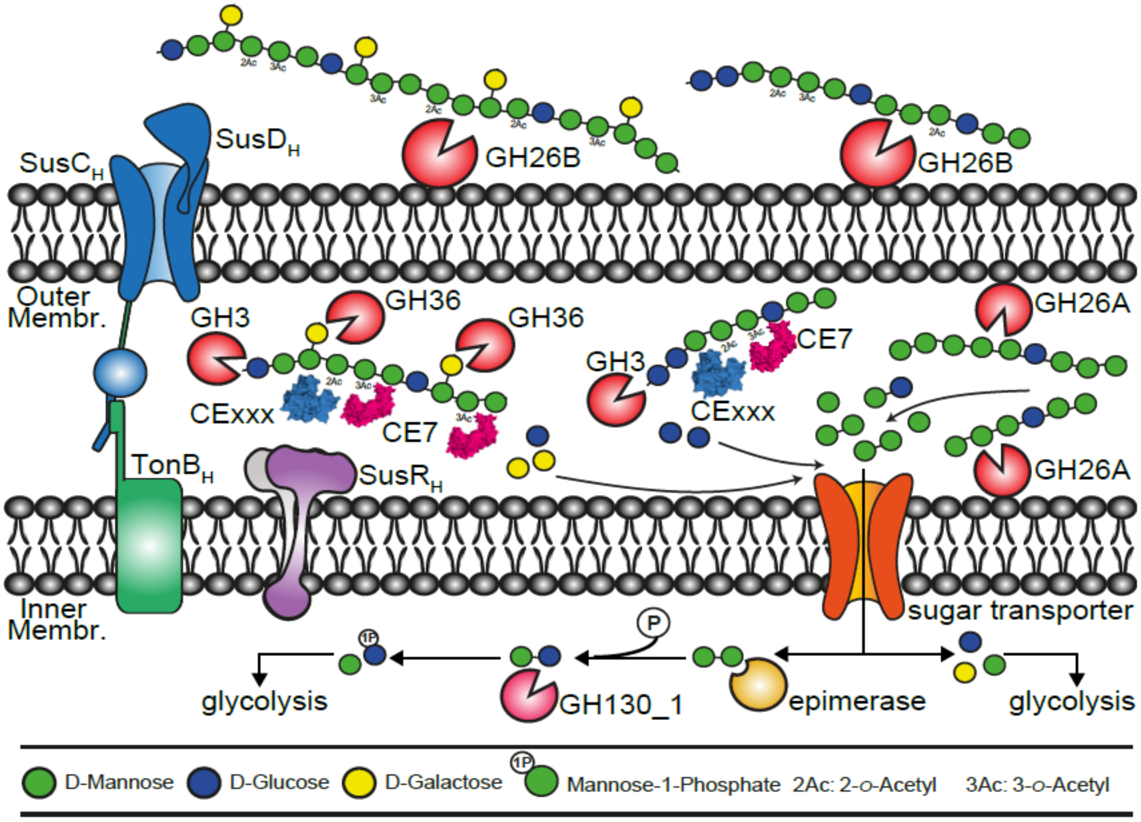


Fig. S8. Proposed β-mannan degradation pathway in *B. cellulosilyticus*. Polymeric β-mannans are depolymerized on the surface of *B. cellulosilyticus* by the endomannanase *Bc*GH26B, and the products are imported into the cell by the SusC-SusD-like transporter proteins. In the periplasm, the galactosyl and acetyl decorations are removed by the α-galactosidase *Bc*GH36 and the acetyl esterases *Bc*CE7 and *Bc*CExxx. The mannanase *Bc*GH26A generates mannobiose or short mannooligosaccharides, and the exo-acting glucosidase *Bc*GH3 removes glucose residues. Mono- and disaccharides are further transported into the cytosol by a sugar transporter. Mannobiose is epimerized to mannosyl-glucose by the epimerase. The phosphorylase *Bc*GH130_1 phosphorolyses mannosyl-glucose to mannose-1-phosphate and glucose. The monosaccharides then enter glycolysis. The specific roles of GH5_2 and GH5_7 are not displayed due to the lack of functional information.
